# A deep learning-based approach for high-throughput hypocotyl phenotyping

**DOI:** 10.1101/651729

**Authors:** Orsolya Dobos, Peter Horvath, Ferenc Nagy, Tivadar Danka, András Viczián

**Affiliations:** Institute of Plant Biology, Biological Research Centre of the Hungarian Academy of Sciences, Temesvári krt. 62, H-6726 Szeged, Hungary; Doctoral School in Biology, Faculty of Science and Informatics, University of Szeged, Szeged, H-6726, Hungary; Institute of Biochemistry, Biological Research Centre of the Hungarian Academy of Sciences, Temesvári krt. 62, H-6726 Szeged, Hungary

**Keywords:** plant phenotyping, Arabidopsis, computer vision, machine learning, deep learning

## Abstract

Hypocotyl length determination is a widely used method to phenotype young seedlings. The measurement itself has been developed from using rulers and millimeter papers to the assessment of digitized images, yet it remained a labour-intensive, monotonous and time consuming procedure. To make high-throughput plant phenotyping possible, we developed a deep learning-based approach to simplify and accelerate this method. Our pipeline does not require a specialized imaging system but works well with low quality images, produced with a simple flatbed scanner or a smartphone camera. Moreover, it is easily adaptable for a diverse range of datasets, not restricted to *Arabidopsis thaliana*. Furthermore, we show that the accuracy of the method reaches human performance. We not only provide the full code at https://github.com/biomag-lab/hypocotyl-UNet, but also give detailed instructions on how the algorithm can be trained with custom data, tailoring it for the requirements and imaging setup of the user.

**One-sentence summary:** A deep learning-based algorithm, providing an adaptable tool for determining hypocotyl or
coleoptile length of different plant species.

## Introduction

Monitoring different aspects of seedling development requires determining certain physical dimensions of the plantlet. Among these, measurement of hypocotyl length is a key phenotypic trait to monitor and quantify different responses. Hypocotyl cells are formed in the embryo and their eventual number set in after only a few cell divisions. During seedling growth, the length of the hypocotyl is determined by no further cell divisions but by the elongation of hypocotyl cells (Gendreau et al., 1997). Hypocotyl growth is regulated by a complex network of external and internal factors. Internal cues are hormones: auxins, ethylene, cytokinins, abscisic acid, gibberellins and brassinosteroids has been shown to be involved in the response (Vandenbussche et al., 2005; Hayashi et al., 2014). Among external cues, gravity not only determines the direction of growth (away from the soil surface) but also affects the hypocotyl elongation (Soga et al., 2018). Our knowledge about how light regulates hypocotyl elongation is much more detailed. Without light, etiolated plants develop elongated hypocotyls, whereas light triggers photomorphogenic development with characteristic, fluence rate dependent inhibition of hypocotyl elongation, which is one of the key feature of the so-called photomorphogenic growth (Fankhauser and Casal, 2004; Arsovski et al., 2012). The role of different light sensing molecules (photoreceptors) has been revealed in this response: phytochrome B (phyB) is the dominant photoreceptor in red (R), phyA in far red (FR) and cryptochrome 1 (cry1) in blue (B) light (Lin et al., 1996; Nagy and Schäfer, 2002). Photomorphogenic ultraviolet B (UV-B) radiation also induces inhibition of hypocotyl elongation (Kim et al., 1998) involving pathways controlled by UV RESISTANCE 8 (UVR8) UV-B receptor (Favory et al., 2009). Fluence rate response curves are used to depict hypocotyl length change over broad light fluences, demonstrating the involvement of specific receptors and their signaling partners in the examined responses. Temperature is the third external cue which have effect on hypocotyl length. It was recently shown how lower temperature shortens hypocotyl length via phyB in light (Jung et al., 2016; Legris et al., 2016; Casal and Qüesta, 2018).

These examples show that hypocotyl length is a seedling phenotypic trait of particular importance. On one hand it indicates the functionality of the examined signaling pathway(s), on the other hand it is relatively easy to measure generating quantified data of the observed response. Thus researchers measure hypocotyl length (i) to compare the effect of different light, hormone etc. treatments, (ii) to analyze the role of signaling components using mutants and overexpressor lines and (iii) to perform different reverse and forward genetic (screening) approaches.

The methodology of the hypocotyl measurement changed over time. In early studies the hypocotyls were simply measured by hand one-by-one using a ruler or a millimeter paper, in many cases rounding the observed value to the nearest millimeter (Köhler, 1978; Liscum and Hangarter, 1991; Pepper et al., 2001; Dieterle et al., 2005). More precise and most widely applied quantification procedure involves the arrangement of seedlings on sticky surfaces or agar plates, subsequent scanning or photographing them and measurement of their hypocotyl length using a digital image processing software (Young et al., 1992; Borevitz and Neff, 2008; Ádám et al., 2013; Das et al., 2016). This approach gives the opportunity to store hypocotyl images and measure them at a later time and even involving other experimenters in the measurement procedure. To speed up this process and reduce the invested work-time, different applications were created to automate the quantification of hypocotyl length (Sangster et al., 2008; Wang et al., 2009; Cole et al., 2011; Spalding and Miller, 2013). These image processing tools have the potential to replace error prone and labour intensive manual image processing and to advance plant phenotyping by enabling high-throughput data analysis. A cornerstone of these algorithms is the plant segmentation, that is, the separation of the plant from the background. This is a difficult task due to the diversity of images, which can be caused for example by different image acquisition setups and conditions. However, good segmentation is a key to downstream analyses such as object boundary detection and midline tracking (Spalding and Miller, 2013). In addition to overall plant segmentation, fully automated identification of different plant subparts such as cotyledons, roots and seedcoats is a significant challenge, which has not been solved reassuringly in the previous efforts. For hypocotyl length measurement, a major difficulty is the localization of hypocotyl-root junction and robust identification of the cotyledon. Tools based on classical segmentation algorithms have trouble identifying these parts for several reasons, for instance high variance in phenotypes, variable imaging conditions or noisy images. Since imaging methods are very different from lab to lab and no gold standard is available, it is essential to provide a data analysis pipeline which works robustly for a diverse set of images.

Up until the recent introduction of deep convolutional neural networks (CNN), this was extremely difficult to achieve. In contrast to classical methods, modern deep convolutional networks are capable to surpass human performance in many image processing tasks, for instance object classification and detection (Geirhos et al., 2018). Instead of relying on hand crafted filters and features, a neural network is able to learn the optimal representation of the data. This makes its performance exceptionally good, and given enough data, a well-trained neural network can generalize for a wide range of datasets. For plant phenotyping, these developments yielded advances for instance in trait identification and genotype/phenotype classification (Pound et al., 2017; Namin et al., 2018).

In this paper, we present a deep learning based approach which is capable to provide quantified seedling phenotype data in a high-throughput manner. Compared to earlier tools, ours is fully automated and achieves human expert accuracy on length measurement tasks for various plant species such as *Arabidopsis thaliana, Sinapis alba* and *Brachypodium distachyon*. The method does not require expensive imaging setups, accurate results can be obtained with a simple flatbed scanner or a smartphone camera. In addition, the measurement itself requires only a few seconds per image, thus reducing the time spent by several orders of magnitude. We provide full access to our algorithm as it is open source and also give detailed instructions how to perform training for customised hypocotyl length determination approaches.

## Results

### The architecture of the algorithm

To extract the length data from images, first we perform segmentation, followed by the skeletonization of the segmented objects to be measured (Fig. 1). In the case of a typical seedling, each image is segmented into three non-overlapping parts: 1) background 2) hypocotyl 3) non-hypocotyl seedling area. (The latter category differs between species, thus different non-hypocotyl parts should be defined accordingly.) Central to our approach is the U-Net deep convolutional neural network for segmentation, which is particularly excellent for finding thin objects. It has been applied with success on various problems such as detecting cell nuclei in microscopic images or identifying subparts of the brain on MRI scans (Ronneberger et al., 2015; Buda et al., 2019). On images for plant phenotyping, in addition to separating the plant from the background, U-Net is able to identify the specific parts of the plants. On a provided image, U-Net applies convolution operations with various filters followed by maximum pooling repeatedly, producing the segmentation masks. The major difference, as opposed to classical image processing algorithms, is that the filters used by the network are not given in advance but learned from the data during the so-called training phase. In this phase, the segmentation masks provided by the expert are shown for the algorithm several times, which is then able to learn how to classify each pixel either as background or a specific plant organ. This training process gives rise to filters which are best suited for the task and data, resulting in an extremely robust and adaptable method.

**Figure 1.**
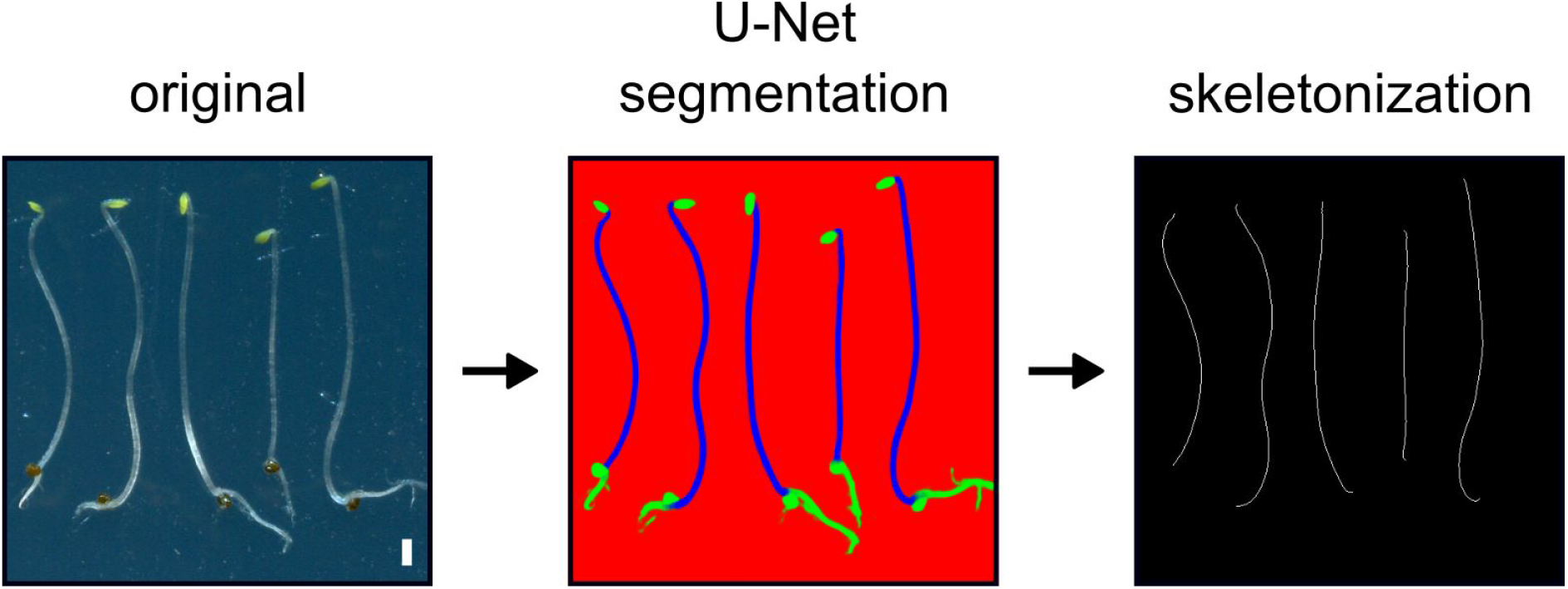
The overview of the method. Arabidopsis seedlings were placed on agar plate surface and scanned resulting in the original image. This image was then processed by the previously trained U-Net algorithm (see *Materials and methods* chapter for details), which determines plant parts what are hypocotyls (marked with blue colour) and also those which are other non-hypocotyl plant parts (depicted by green colour). This step is called segmentation. During the next step, the algorithm determines a 1-pixel-wide line in the middle of the segmented hypocotyls. This procedure is called skeletonization and the number of pixels consisting of the 1-pixel-wide lines is proportional with the hypocotyl length. White scale bar represents 1 mm.

After the specific plant parts are segmented and identified, the binary images of the hypocotyls are skeletonized (Lee et al., 1994). Skeletonization is the reduction of binary shapes to 1 pixel wide representations, a curve in the case of hypocotyls. This operation allows the length measurement of spatial objects. On the skeleton image, components representing hypocotyls were measured by calculating the number of pixels for each identified object and then converted from pixel unit to *mm*. Pixel to *mm* calculations were performed by either scaling directly with the DPI (dots-per-inch) value of the image or using a reference object on each image. After the measurement, very small objects, which are most likely due to segmentation errors, are filtered out. Finally, the obtained results are exported as a *csv* file, ready for downstream analysis.

### The choice of the convolutional network architecture

In general, a convolutional neural network repeatedly performs convolutional, pooling and in some instances, batch normalizing operations, eventually extracting a feature-level representation of the image. This is called encoding. During this part, information is compressed and can be lost during the pooling steps. For tasks such as image classification, this is not a problem (Pound et al., 2017). However, for semantic segmentation tasks, the network is required to reconstruct the pixel-level segmentation mask, which is achieved by upsampling the feature-level representation. In this decoding step, the information lost during encoding cannot be recovered and will result in suboptimal results for small or thin objects, such as hypocotyls in our case. This problem was solved with the introduction of U-Net (Ronneberger et al., 2015), originally created to find cells in microscopy images, where the cells can grow on each other, having only a thin (occasionally 1-2 pixel wide) region separating them. This is achieved by storing the intermediate feature-level representations before each pooling in the encoding step, then feeding this data to the corresponding upsampling layer. Ever since its inception, U-Net has become a state of the art architecture for semantic segmentation. Because of its performance on small or thin objects, this choice of architecture was ideal for our purposes. To add a regulazing term and accelerate training speed, we have added batch normalizing layers after convolutional blocks (Ioffe and Szegedy, 2015).

### Phenotypic analysis of Arabidopsis seedlings

Determining hypocotyl length of Arabidopsis seedlings is a key phenotyping procedure in myriads of studies thus it was obvious to test our algorithm on this model plant first. We simply grew seedlings on wet filter papers under different fluences of monochromatic light, laid them on agar plates, scanned them and then used these images to train the algorithm. Altogether we annotated about 2500 hypocotyls and corresponding non-hypocotyl plant parts during this procedure. To test the trained algorithm, we grew seedlings under different fluences of monochromatic red light as a routine treatment for phytochrome studies. Fig. 2A shows, how the algorithm recognized long and short hypocotyls belonging to those plants which grow under low or high fluences of light, respectively. The fluence rate graph plotting the measured hypocotyl length values demonstrate that the algorithm determined values similar to the human experimenters (Fig. 2B). To further test the versatility of the algorithm we analyzed hypocotyls of seedlings grown in far-red and blue light when the inhibition of hypocotyl elongation is mediated by phyA and cryptochrome photoreceptors, respectively. Additionally we analyzed also etiolated seedlings grown in darkness which are used as important controls in photobiological studies. We found that the performance of the algorithm is comparable to humans under these conditions and the measurement works well even in case of the pale, almost colourless etiolated seedlings (Supplemental Figs. S1-S3). It was tempting to further examine seedlings which have completely different body architecture. For this purpose, we grew plantlets on plant medium containing sugar with white light illumination. These seedlings have thick hypocotyls, fully developed and opened green cotyledons and long roots. Our results show that the algorithm is capable to measure the hypocotyls of seedlings grown under light/dark cycles or under continuous white light supplemented with or without photomorphogenic (non-damaging) UV-B irradiation Supplemental Fig. S3.

**Figure 2.**
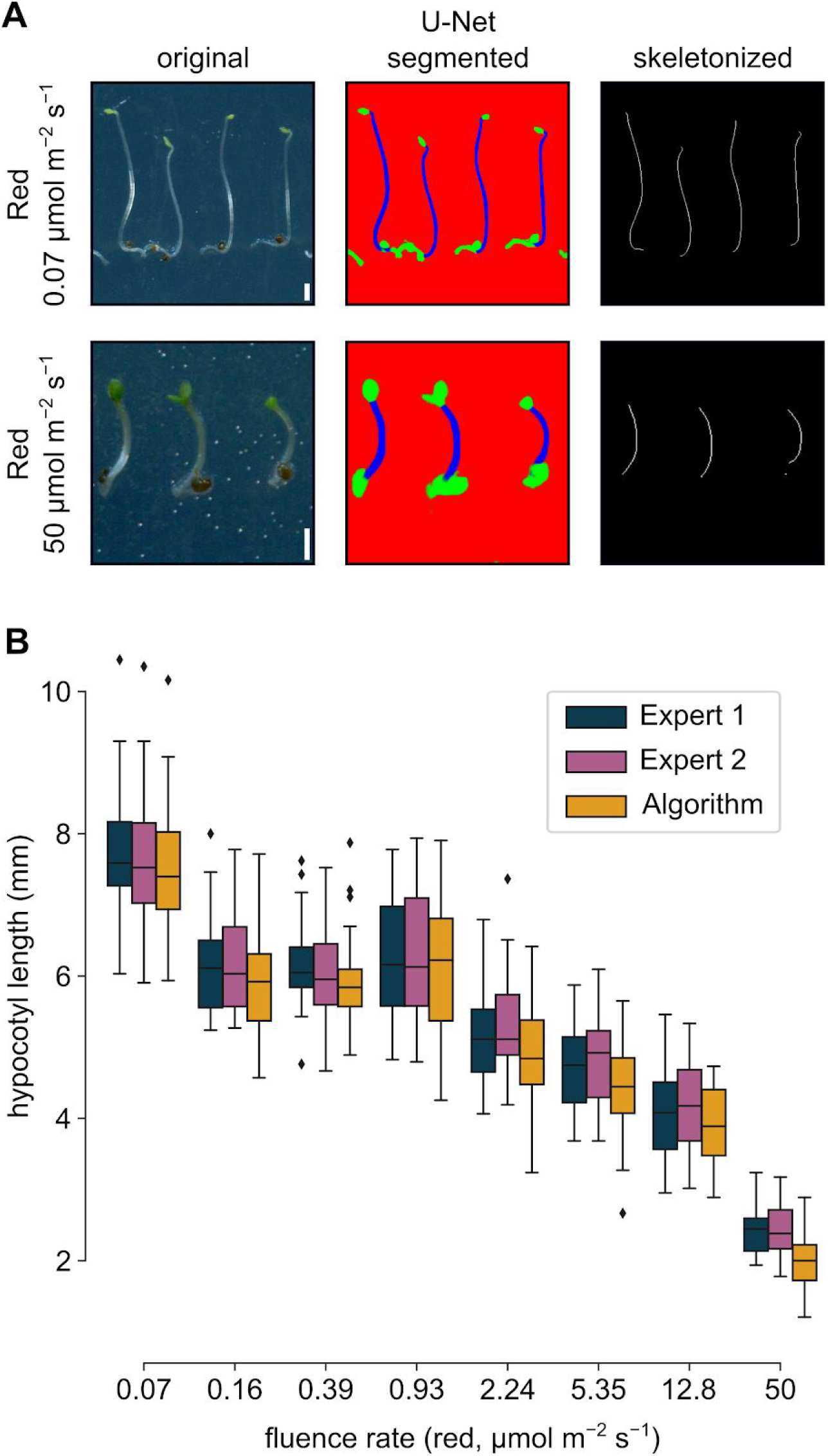
Hypocotyl measurement of red light-grown Arabidopsis seedlings. (a) Arabidopsis seedlings were grown on wet filter papers in red light for 4 days, placed on agar plate and scanned. A close-up image shows a few seedlings grown under high or low fluences of red light, the U-Net segmented and skeletonized image generated from the original by our algorithm. Scale bars represent 1 mm. (b) Fluence rate graph presenting the hypocotyl length values determined by the algorithm and two human experimenters. This box-and-whisker diagram shows the distribution of seedling hypocotyl length, where boxes depict the quartiles, whiskers extend to show the rest of the distribution. Black diamonds represent outliers. Sample number at every data point is n=30.

### Application of the algorithm on different plant species

To test the usability of our algorithm on other species besides Arabidopsis, we choose mustard (*Sinapis alba*) and stiff brome (*Brachypodium distachyon*). Mustard was a widely used experimental object a few decades ago to examine the dependency of hypocotyl elongation on different irradiation protocols. These works revealed the basic mechanisms of phytochrome action many years before identifying the involved molecular pathways or even the genes coding the photoreceptors (Schopfer and Oelze-Karow, 1971; Wildermann et al., 1978a; Wildermann et al., 1978b). A recent study demonstrated that determining the hypocotyl elongation of mustard seedlings is still in use as a phenotypic marker to monitor hormonal changes under different irradiation conditions (Procko et al., 2014).

The mustard plantlets were grown on agar plates under constant white light for 4 days. These seedlings were too bulky to scan them with a flatbed scanner like we did with the Arabidopsis seedlings. For this reason, images were taken with a smartphone. We used these images to train our algorithm to identify pixels belonging to mustard hypocotyls and to determine hypocotyl length. During the training phase we annotated about 250 hypocotyls and corresponding non-hypocotyl plant parts before performing the presented measurement. Fig. 3 demonstrates that this low number of seedlings were enough to train the algorithm to determine hypocotyl length with high accuracy, which is comparable to the performance of the human experts.

**Figure 3.**
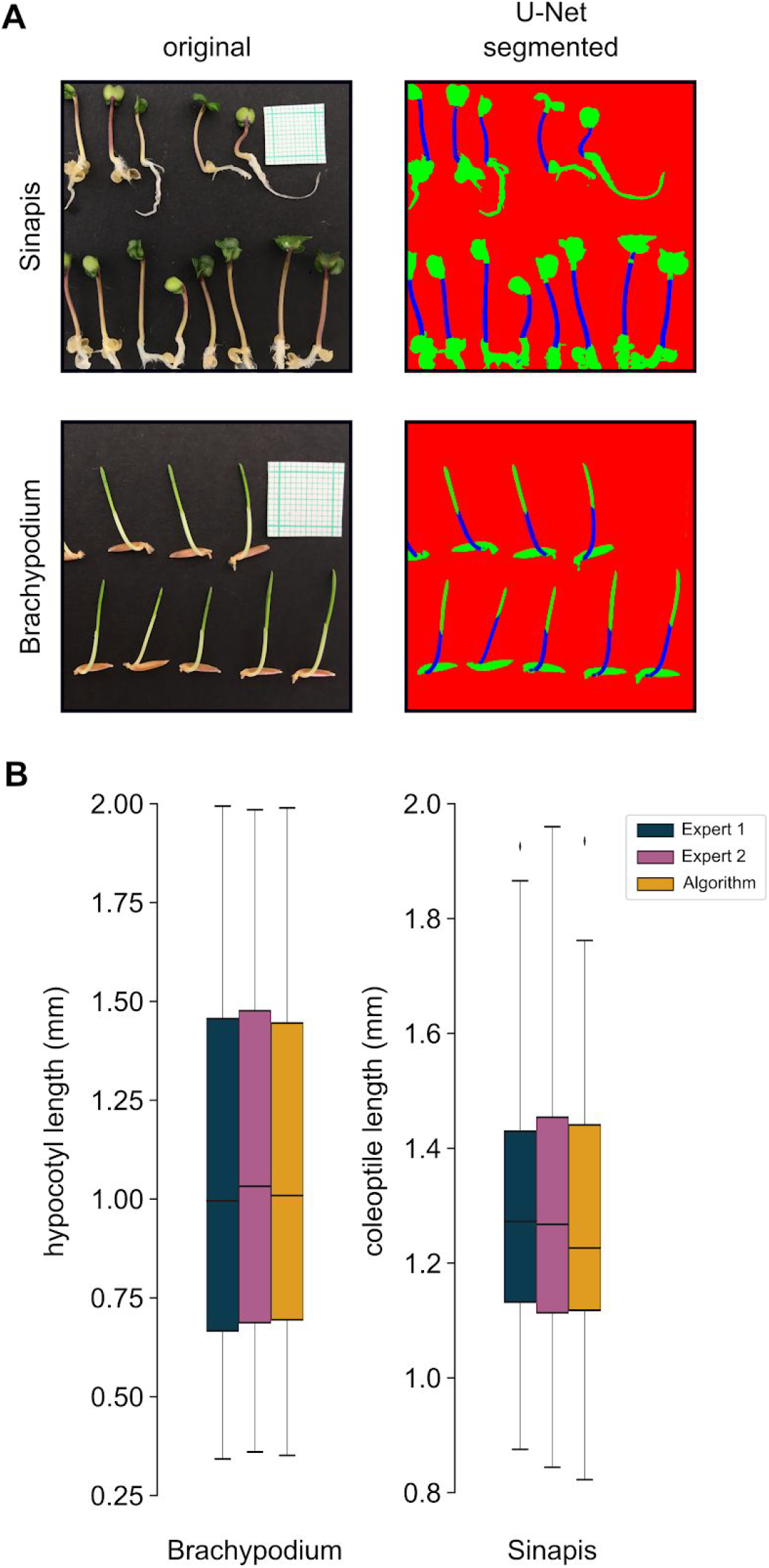
Mustard hypocotyl and Brachypodium coleoptile measurements by the algorithm. (a) Original images of light-grown mustard and Brachypodium plantlets (right side). Image panels at the left side depict the segmentation made by the algorithm. The original images also contain the millimeter paper for size scaling purposes. (b) Box-and-whisker diagrams show the hypocotyl length values determined by the U-net algorithm and two human experts. Boxes depict the quartiles, whiskers extend to show the rest of the distribution, whereas black diamonds represent outliers. Sample number in case of the mustard seedlings is n=91, and in case of the Brachypodium plantlets it is n=75.

We further tested the versatility of the algorithm by analyzing monocotyledonous plants. In monocots, the coleoptile growth is a widely used phenotypic trait instead of the more difficulty observable hypocotyl. We chose stiff brome (*Brachypodium distachyon*), which is a small-sized plant, having compact and sequenced genome (International Brachypodium Initiative, 2010) and an existing transformation system (Alves et al., 2009). These make it an ideal grass model species with emerging importance (Scholthof et al., 2018). We grew the Brachypodium plants under red light or in darkness for 4 days and took photos of them with a smartphone camera. In this case we used 8 images containing about 100 plants to train the algorithm. Fig. 3 shows how the algorithm processed the images and that how it measured coleoptile length on the test images. The obtained values do not differ from those what the human experts measured, demonstrating the usability of the algorithm to analyze Brachypodium coleptiles.

### Accuracy of the algorithm

To quantitatively assess the performance of our algorithm, we decided to compare the obtained results to the performance of humans. Each measurement was repeated by two human experimenters. For each seedling identified by the algorithm, we calculated measurement accuracy by matching the seedling to the ground truth data provided by the experts (Fig. 4). Matching was done by scanning through the hypocotyls measured by the experts and comparing bounding boxes of the identified seedling and the ground truth. To accept a plant as a match, we required at least 10% overlap of the bounding boxes between the identified object and the plant. (Since the seedlings were placed apart from each other on the plates, the possibility of a false matching was minimal.) This provided a way to calculate the false positive (FP) and true positive (TP) ratio, and in the case of a match, the accuracy of the length measurement was assessed as well, which is defined as the relative error between the ground truth value and the algorithm’s result.

**Figure 4.**
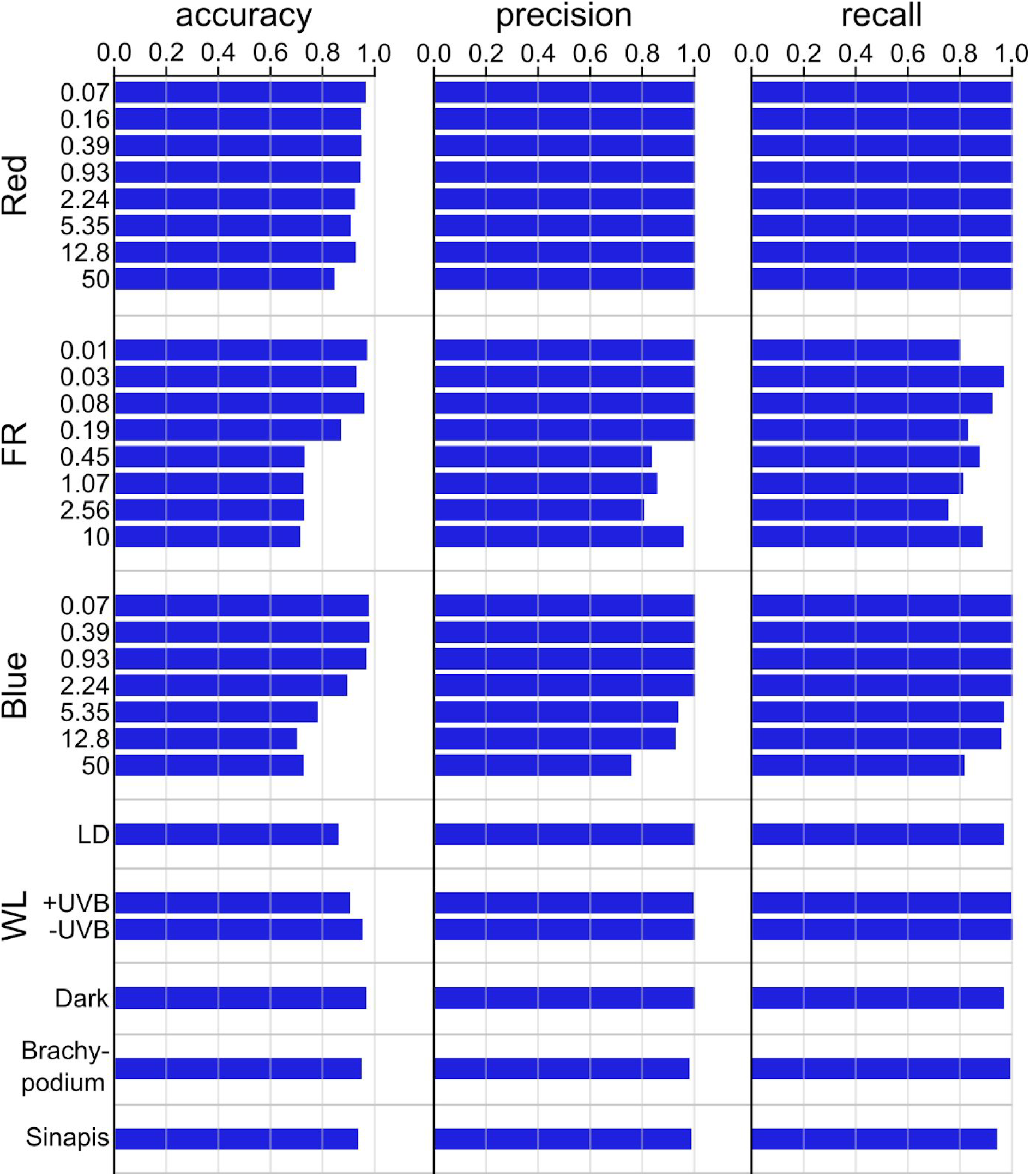
Accuracy, recall and precision metrics for the algorithm for each light condition. Further analysis of the data is presented in Figs 2, 3 and Supplemental Figures S1-S3. *Accuracy* was calculated by first matching the plants identified by the algorithm to the ground truth given by the experts, and if a match was found, the relative accuracy of the measurements were calculated. (A match is required to have at least 10% overlap between the bounding boxes of the objects.) The *precision* of the algorithm is defined as TP/(TP + FP), where TP and FP denotes the number of true and false positives, respectively. That is, a high precision implies that the majority of identified objects are indeed plants, not opposed to false detections. Finally, *recall* is given by TP/(TP + FN), where FN is the number of false negatives. The higher the recall, the more plants were identified by the algorithm.

For a more detailed view on the detection performance, we also calculated the precision and the recall values. Precision is defined by TP/(TP + FP), where TP and FP are the number of true positives and false positives respectively, whereas recall can be calculated by TP/(TP + FN), with FN denoting the number of false negatives. We calculated accuracy, recall and precision individually to each plant, compared to the measurement of each expert, then averaged together. For all of our metrics, a higher value implies a better result (Fig. 4). To put this in perspective, a high precision means that most identified objects are indeed plants (as opposed to for instance segmentation errors), whereas a high recall means that most plants were indeed detected in the image. In general, there is a tradeoff between recall and precision, which is controlled by the strictness of our criteria to accept a match. A too loose criteria leads to an abundance of false detections resulting in potentially high recall but very low precision. On the other hand, an excessively strict criteria would result in a high false negative rate, leading to low recall and potentially high precision. Thus, the combination of recall and precision together provides a good description on the performance of the algorithm.

To obtain further data to characterize the hypocotyl measurement, as the method itself, both human experimenters measured each plant once more, having one month between their two measurements. Using these repeated measurements, we calculated the intra-expert accuracy exactly as we outlined above, using the two measurements provided by the same expert. The inter-expert accuracy was calculated using the first measurement of both experts. We found that the algorithm performs exceptionally well on plants with long hypocotyls, but with slightly lower reliability in case of the very short seedlings grown under strong FR or B light. It is also important to note that the performance of the humans is also poorer when analyzing these plantlets both in case of intra- or inter-expert comparisons (Fig. 5).

**Figure 5.**
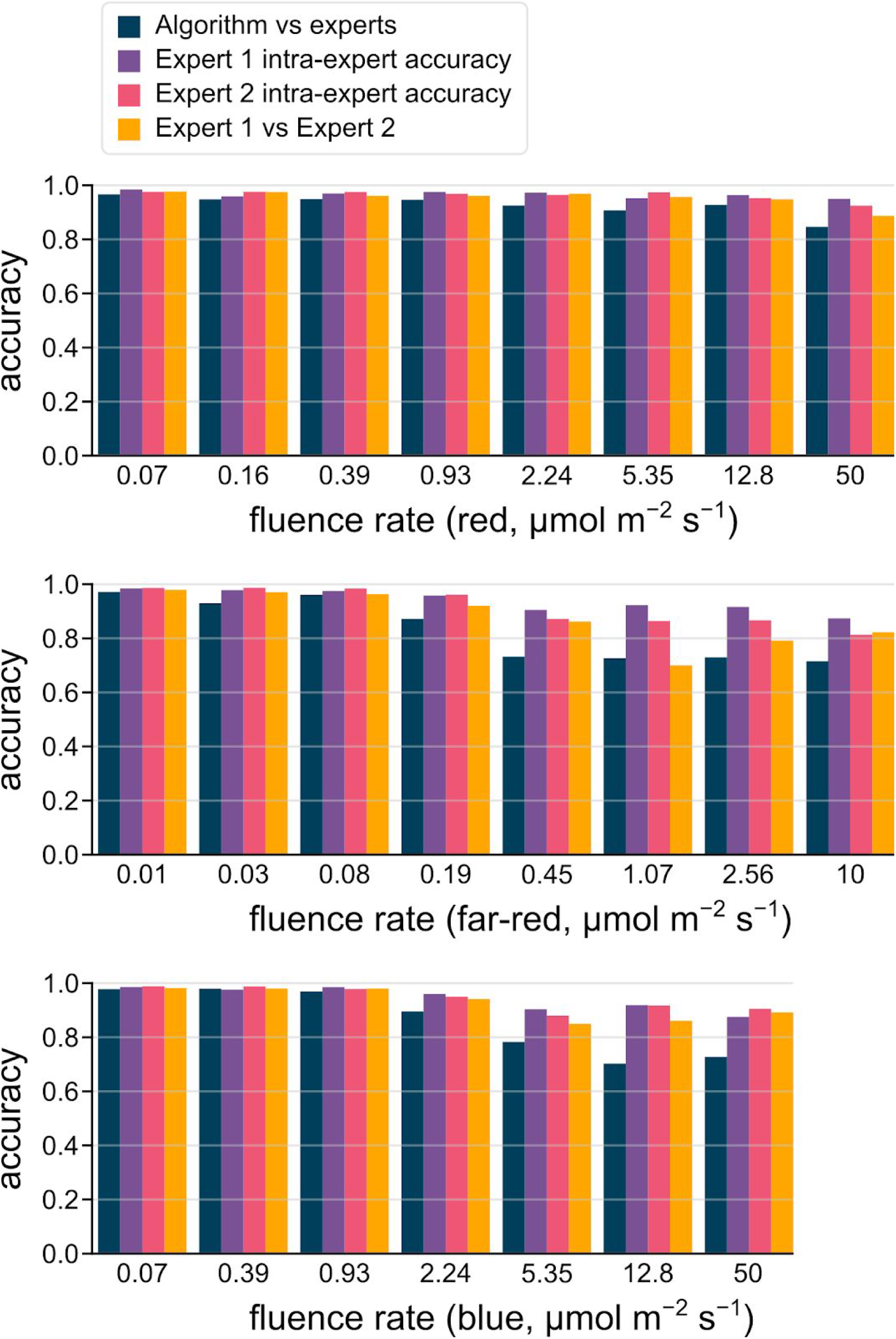
Intra- and inter-expert accuracies vs the algorithm. *Intra-expert accuracy* was calculated by averaging the accuracies between the two measurements from the same expert. *Inter-expert accuracy* (Expert 1 vs Expert 2) was determined by comparing the first measurements of human experts. For comparison, the accuracy of the algorithm is also presented.

## Discussion

### Usability of the method

Hypocotyl growth is controlled by the interplay of different external and internal cues, many of them with reciprocal effects. It follows, that hypocotyl length is used (i) to characterise activity of numerous signalling pathways, including those controlled by light, hormones, temperature and gravity and that (ii) determination of hypocotyl length is a widely used basic seedling phenotyping assay. Here we report the development of a deep learning-based algorithm to simplify this measurement and save valuable time for the experimenter. There have been computer-based tools published earlier, but our work presents a truly novel approach by demonstrating the suitability of deep learning for quantitative plant phenotyping. This method is applicable to a diverse set of image-based phenotyping problems, not restricted to hypocotyl measurement. Our method uses the U-Net convolutional neural network architecture for segmentation and it is able to identify not only hypocotyls, but also roots and cotyledons with previously unprecedented detail. To demonstrate the power of the algorithm, we have shown how it performs on other dicot or monocot seedlings. The method possess several advantages: (i) no image preprocessing is needed; (ii) the algorithm is capable to handle low quality images, i.e. ones made with a simple smartphone camera; (iii) very different imaging conditions and (iv) its performance is matching human accuracy. Moreover, the whole measurement pipeline is fully automated, it does not require manual intervention at all. This decreases the execution time with several orders of magnitude: while the expert spends 45 minutes on average to manually measure a complete image and record the data, our method performs the same task under a minute. With this speedup, high-throughput assays (testing numerous lines, phenotype-based screenings, etc) are enabled for a wide array of questions.

### Assessing our results

To assess the performance of our algorithm, first we focused on Arabidopsis, being the most widely used model plant. Our algorithm performed quite well on seedlings with various body architectures. We tested it on seedlings having short or long, thick or thin hypocotyls; opened or unopened cotyledons with different thickness, size and colour; roots with different length, shape and thickness. The accuracy, the precision and recall values, compared to the results of a human experimenter indicate that the algorithm is suitable to replace manual measurements for a wide array of scenarios (Figs. 4 and 5). Our data also shows that under specific circumstances, when the plants are short (under strong FR and blue light), the accuracy of the algorithm is slightly lower compared to human experimenters. The reasons are quite diverse.

(i) The accuracy value is heavily affected by the absolute size of the plant. For example, a 5 pixel error on a 100-pixel-long plant has 0.95 accuracy, whereas on a 20-pixel-sized one, the same absolute error yields 0.75 accuracy. (In our images, a typical hypocotyl length of a seedling grown under high light intensities is appeared as only around 20 pixels.)

(ii) In case of short and thick hypocotyls, human experts cannot position their region of interest (ROI) at the middle of the hypocotyl. In this case the skeletonization can be different from the human ROI placement.

(iii) Misplaced seedlings (hypocotyls touching each other, roots laying over the hypocotyl, etc) or image problems (reflecting plastic plate edges, scratches of the agar surface) disturb the segmentation process, but with less extent as the human experts. These issues can be corrected manually on the generated data and also a certain carefulness is required during seeding placement onto the agar before the scanning. Another potential source of inaccuracy is the skeletonization of the segmented hypocotyls. Especially for more complicated shapes and cusps, the skeletons may have small additional branches or may not be simply connected at all, which can distort the length measurements.

(iv) Especially in the case of seedlings having short and thick hypocotyls, it is not obvious how to define the border between the hypocotyl and the root. For that, images with higher magnification (i.e microscopy) should be obtained, which is not manageable when working with high number of seedlings (Fahn, 1990). This problem is a general caveat of the method: the observable morphological traits at the resolution of the scanned images are not sufficient sometimes to mark precisely where does the hypocotyl ends and the root begins. Taken together, the inaccuracy generated in these ways, is an inevitable component of hypocotyl measurement leading to the errors, not only in case of the algorithm, but also in case of measurements made by humans (Figs. 4 and 5). Similarly to the algorithm, the expert accuracy also decays as when working with small seedlings. However, under these conditions, the expert performance is 10-20% better than the algorithm, although at some points the inter-expert (experts compared to each other) accuracy is not better than the accuracy of the algorithm compared to the experts (Fig. 5). Notwithstanding this, we speculate that by training the algorithm exclusively on small plants, the performance could be on par with the experts, although this version of the algorithm could provide good results for small plants only. Conclusively, without having solid ground truth data, the training of the algorithm is unavoidably impaired. During the training procedure we annotated about 2500 Arabidopsis hypocotyls, whereas annotating approximately 250 mustard seedlings and about 100 Brachypodium coleoptiles were sufficient to reach similar recognition metric parameters. These data indicate that Arabidopsis is a ‘difficult’ experimental object in terms of hypocotyl measurement, although we must note that our algorithm trained for Arabidopsis is suitable to analyse seedlings with much diverse plant architecture, whereas in case of the two other species we worked with plantlets grown under only certain conditions.

### Future outlook

In recent years, the introduction of deep learning and convolutional neural networks revolutionized computer vision-based research, making the automation of various tasks and precise high-throughput phenotyping available for many disciplines. In plant biology, several advances have been made with these methods regarding qualitative phenotyping (Pound et al., 2017; Namin et al., 2018; Pineda et al., 2018; Singh et al., 2018; Ramcharan et al., 2019). With these tools however, quantitative phenotypic traits can also be assessed as we demonstrated in this work. The presented segmentation pipeline is not only applicable to length measurements, but in principle it can also be used to measure other parameters such as cotyledon area, hypocotyl hook opening, angle of cotyledons, etc. With the elimination of manual measurements, the current bottleneck in the phenotyping workflow is the ordered laying of the plantlets onto agar plates with special care to avoid overlaps between the plants. This labour-intensive step can be eliminated using object detection frameworks such as Mask-RCNN (He et al., 2017), however, at present these may cause additional segmentation errors, thus reducing accuracy.

While different technical aspects still remain to overcome, we believe that increasing application and improvement of convolutional neural networks for image-based analysis of plants are laying down the foundation for the next generation of plant phenotyping tools.

## Materials and methods

### Code and data availability

The algorithm was implemented in Python, where the PyTorch framework was used for deep learning and scikit-image library was used for image processing (van der Walt et al., 2014). The code is fully open source and available at GitHub (https://github.com/biomag-lab/hypocotyl-UNet). Images used for training are also available at https://www.kaggle.com/tivadardanka/plant-segmentation.

### Image acquisition and data preparation

Arabidopsis seedlings were laid manually onto the surface of 1% agar plates. During scanning a black paper sheet was used as reflective document mat. The scanning was done by EPSON PERFECTION V30 scanner, at 800 dpi and 24-bit color setting, pictures were saved as. tif or. jpg. After acquisition, the hypocotyl, cotyledon, seedcoats and roots were annotated using Fiji (Schindelin et al., 2012), which was used to create the mask for training the segmentation algorithm. Before training, the images were padded by mirroring a 256 pixel wide strip next to the border. The padded images were cropped up to non-overlapping pieces with 800×800 resolution which were used to train the neural network. During training, 10% of the images were held out for validation purposes.

### Training the neural network

To train the U-Net convolutional neural network for plant segmentation, we have annotated about 2500 Arabidopsis hypocotyls, 250 Sinapis seedlings and 100 Brachypodium plantlets. For each of the plant species, a different U-Net model was trained. More details on the U-Net architecture can be found in (Ronneberger et al., 2015). As additional regularization, batch normalization layers were used after the convolutional blocks, which was shown to be highly effective for such CNN architectures (Ioffe and Szegedy, 2015). During training, the smooth Dice coefficient loss was used, introduced by (Milletari et al., 2016; Sudre et al., 2017). The model was trained to classify each pixel as 1) background 2) hypocotyl (or coleoptile in case of Brachypodium) 3) plant parts not included in the measurement (root, cotyledon, seedcoat, etc.). To assure that the plant parts are precisely segmented, their corresponding term in the loss function was weighted fivefold compared to the background. Training was run for 1000 epochs with initial learning rate 1e-4, which was consequently decreased during training to 1e-5, 1e-6 and 1e-7 after epochs 200, 600 and 900. The algorithm was trained using a single nVidia Titan XP GPU. For optimization, the Adam optimizer was used (Kingma and Ba, 2014). To prevent overfitting, batch normalization and image augmentation was used. The augmentation transform was composed as a series of random 512×512 crops, affine transforms with flips and a color jitter transform. The detailed procedure of reproducing our workflow is described as an instructional help document in the Supplemental Method S1-S2. The annotated images used for training can be found at https://www.kaggle.com/tivadardanka/plant-segmentation. All presented hypocotyl and coleoptile length data, was measured on images which were not involved in the training procedure.

### Plant growth conditions and light treatments

*Arabidopsis thaliana* (Columbia 0 (Col-0) ecotype) seeds were sown on 4 layers of wet filter paper and were kept at 4 °C for 3 days. In order to promote homogeneous germination, plates were exposed to 70-100 μmol m^−2^ s^−1^ white light for 8 h (LUMILUX XT T8 L 36 W/865 fluorescent tubes, Osram), followed by exposition to continuous red (λ_max_= 660nm), far-red (λ_max_= 735 nm) or blue (λ_max_= 470 nm) light for 4 days at 22 °C (SNAP-LITE LED light sources, Quantum Devices, WI, USA). Plates containing dark-grown seedlings seedlings plates were wrapped in aluminum foil and kept in dark for 4 days at 22°C.

Seeds sown on ½ Murashige and Skoog (MS, Sigma-Aldrich) medium containing 1% sucrose and 0.8% agar were surface sterilised and kept at 4 °C for 3 days. Seedlings were grown under 12 h white light (80 μmol m^−2^ s^−1^)/ 12 h dark photocycles at 22 °C in a growth chamber (MLR-350H, SANYO, Gallenkamp, UK) for 7 days. Alternatively, after 3 days, plates were placed under continuous white light (PHILIPS TL-D 18 W/33-640 tubes, 1.5 *μ*mol m^−2^ s^−1^) supplemented with UV-B (PHILIPS ULTRAVIOLET-B TL20W/01RS tubes, 10 *μ*mol m^−2^ s^−1^)) for 4 days at 22 °C. The seedlings were covered with transmission cut-off filters (WG series, Schott), using the WG305 filter for UV-B-treated seedlings (+UV-B), and the WG385 filter for the control (-UV-B) seedlings as providing half maximal transmission at 305 or 385 nmm, respectively (Bernula et al., 2017).

*Brachypodium distachion* (Bd21) seeds were sown on 1% agar and kept at 4 °C for 5 days and were treated with 24 h white light (130 μmol m^−2^ s^−1^) to induce even germination. Seedlings were grown either in darkness or under 50 μmol m^−2^ s^−1^ red light for 4 days. Subsequently, they were placed on a matte black cardboard sheet and illuminated with even diffused light. Images of the seedlings were taken with a smartphone (iPhone SE, Apple) using the default settings of the camera. Every image contained a millimeter paper for scaling.

Mustard (*Sinapis alba* L.) seeds were sown on 1% agar and kept at 4 °C for 5 days. Seedlings were grown under 130 μmol m^−2^ s^−1^ white light at 22 °C for 4 days. Seedlings were photographed as described for Brachypodium plants.

## Acknowledgments

We thank for Dr. János Györgyei providing the Brachypodium seeds and giving advices on seedling propagation. The work was supported by grants from the Economic Development and Innovation Operative Program (GINOP-2.3.2-15-2016-00001, GINOP-2.3.2-15-2016-00015 and GINOP-2.3.2-15-2016-00026). T.D. and P.H.

acknowledge support from the HAS-LENDULET-BIOMAG and from the European Union and the European Regional Development Fund.

## References

Ádám É, Kircher S, Liu P, Mérai Z, González-Schain N, Hörner M, Viczián A, Monte E, Sharrock RA, Schäfer E, et al(2013) Comparative functional analysis of full-length and N-terminal fragments of phytochrome C, D and E in red light-induced signaling. New Phytol 200: 86–96

Alves SC, Worland B, Thole V, Snape JW, Bevan MW, Vain P (2009) A protocol for Agrobacterium-mediated transformation of Brachypodium distachyon community standard line Bd21. Nat Protoc 4: 638–649

Arsovski AA, Galstyan A, Guseman JM, Nemhauser JL (2012) Photomorphogenesis. Arabidopsis Book 10: e0147

Bernula P, Crocco CD, Arongaus AB, Ulm R, Nagy F, Viczián A (2017) Expression of the UVR8 photoreceptor in different tissues reveals tissue-autonomous features of UV-B signalling: UVR8 signalling in different tissues. Plant Cell Environ 40: 1104–1114

Borevitz J, Neff M (2008) Phenotypic analysis of Arabidopsis mutants: hypocotyl length. CSH Protoc 2008: db.prot4962

Buda M, Saha A, Mazurowski MA (2019) Association of genomic subtypes of lower-grade gliomas with shape features automatically extracted by a deep learning algorithm. Comput Biol Med 109: 218–225

Casal JJ, Qüesta JI (2018) Light and temperature cues: multitasking receptors and transcriptional integrators. New Phytol 217: 1029–1034

Cole B, Kay SA, Chory J (2011) Automated analysis of hypocotyl growth dynamics during shade avoidance in Arabidopsis. Plant J 65: 991–1000

Das D, St Onge KR, Voesenek LACJ, Pierik R, Sasidharan R (2016) Ethylene- and Shade-Induced Hypocotyl Elongation Share Transcriptome Patterns and Functional Regulators. Plant Physiol 172: 718–733

Dieterle M, Thomann A, Renou J-P, Parmentier Y, Cognat V, Lemonnier G, Müller R, Shen W-H, Kretsch T, Genschik P (2005) Molecular and functional characterization of Arabidopsis Cullin 3A. Plant J 41: 386–399

Fahn A (1990) Plant anatomy. Pergamon

Fankhauser C, Casal JJ (2004) Phenotypic characterization of a photomorphogenic mutant. Plant J 39: 747–760

Favory J-J, Stec A, Gruber H, Rizzini L, Oravecz A, Funk M, Albert A, Cloix C, Jenkins GI, Oakeley EJ, et al(2009) Interaction of COP1 and UVR8 regulates UV-B-induced photomorphogenesis and stress acclimation in Arabidopsis. EMBO J 28: 591–601

Geirhos R, Temme CRM, Rauber J, Schütt HH, Bethge M, Wichmann FA (2018) Generalisation in humans and deep neural networks. In S Bengio, H Wallach, H Larochelle, K Grauman, N Cesa-Bianchi, R Garnett, eds, Advances in Neural Information Processing Systems 31. Curran Associates, Inc., pp 7549–7561

Gendreau E, Traas J, Desnos T, Grandjean O, Caboche M, Höfte H (1997) Cellular basis of hypocotyl growth in Arabidopsis thaliana. Plant Physiol 114: 295–305

Hayashi Y, Takahashi K, Inoue S-I, Kinoshita T (2014) Abscisic acid suppresses hypocotyl elongation by dephosphorylating plasma membrane H(+)-ATPase in Arabidopsis thaliana. Plant Cell Physiol 55: 845–853

He K, Gkioxari G, Dollar P, Girshick R (2017) Mask R-CNN. 2017 IEEE International Conference on Computer Vision (ICCV). IEEE, pp 2980–2988

International Brachypodium Initiative (2010) Genome sequencing and analysis of the model grass Brachypodium distachyon. Nature 463: 763–768

Ioffe S, Szegedy C (2015) Batch normalization: accelerating deep network training by reducing internal covariate shift. Proceedings of the 32nd International Conference on International Conference on Machine Learning - Volume 37. JMLR.org, pp 448–456

Jung J-H, Domijan M, Klose C, Biswas S, Ezer D, Gao M, Khattak AK, Box MS, Charoensawan V, Cortijo S, et al(2016) Phytochromes function as thermosensors in Arabidopsis. Science 354: 886–889

Kim BC, Tennessen DJ, Last RL (1998) UV-B-induced photomorphogenesis in Arabidopsis thaliana. Plant J 15: 667–674

Kingma DP, Ba J (2014) Adam: A Method for Stochastic Optimization. arXiv [cs.LG]

Köhler D (1978) The Course of Ortho-Geotropic Reactions of Shoots. Zeitschrift für Pflanzenphysiologie 87: 463–467

Lee TC, Kashyap RL, Chu CN (1994) Building Skeleton Models via 3-D Medial Surface Axis Thinning Algorithms. CVGIP: Graphical Models and Image Processing 56: 462–478

Legris M, Klose C, Burgie ES, Rojas CCR, Neme M, Hiltbrunner A, Wigge PA, Schäfer E, Vierstra RD, Casal JJ (2016) Phytochrome B integrates light and temperature signals in Arabidopsis. Science 354: 897–900

Lin C, Ahmad M, Cashmore AR (1996) Arabidopsis cryptochrome 1 is a soluble protein mediating blue light-dependent regulation of plant growth and development. Plant J 10: 893–902

Liscum E, Hangarter RP (1991) Arabidopsis Mutants Lacking Blue Light-Dependent Inhibition of Hypocotyl Elongation. Plant Cell 3: 685–694

Milletari F, Navab N, Ahmadi S-A (2016) V-Net: Fully Convolutional Neural Networks for Volumetric Medical Image Segmentation. 2016 Fourth International Conference on 3D Vision (3DV). doi: 10.1109/3dv.2016.79

Nagy F, Schäfer E (2002) Phytochromes control photomorphogenesis by differentially regulated, interacting signaling pathways in higher plants. Annu Rev Plant Biol 53: 329–355

Namin ST, Esmaeilzadeh M, Najafi M, Brown TB, Borevitz JO (2018) Deep phenotyping: deep learning for temporal phenotype/genotype classification. Plant Methods 14: 66

Pepper AE, Seong-Kim M, Hebst SM, Ivey KN, Kwak SJ, Broyles DE (2001) shl, a New set of Arabidopsis mutants with exaggerated developmental responses to available red, far-red, and blue light. Plant Physiol 127: 295–304

Pineda M, Pérez-Bueno ML, Barón M (2018) Detection of Bacterial Infection in Melon Plants by Classification Methods Based on Imaging Data. Front Plant Sci 9: 164

Pound MP, Atkinson JA, Townsend AJ, Wilson MH, Griffiths M, Jackson AS, Bulat A, Tzimiropoulos G, Wells DM, Murchie EH, et al(2017) Deep machine learning provides state-of-the-art performance in image-based plant phenotyping. GigaScience. doi: 10.1093/gigascience/gix083

Procko C, Crenshaw CM, Ljung K, Noel JP, Chory J (2014) Cotyledon-Generated Auxin Is Required for Shade-Induced Hypocotyl Growth in Brassica rapa. Plant Physiol 165: 1285–1301

Ramcharan A, McCloskey P, Baranowski K, Mbilinyi N, Mrisho L, Ndalahwa M, Legg J, Hughes DP (2019) A Mobile-Based Deep Learning Model for Cassava Disease Diagnosis. Front Plant Sci 10: 272

Ronneberger O, Fischer P, Brox T (2015) U-Net: Convolutional Networks for Biomedical Image Segmentation. Medical Image Computing and Computer-Assisted Intervention – MICCAI 2015. Springer International Publishing, pp 234–241

Sangster TA, Salathia N, Undurraga S, Milo R, Schellenberg K, Lindquist S, Queitsch C (2008) HSP90 affects the expression of genetic variation and developmental stability in quantitative traits. Proc Natl Acad Sci U S A 105: 2963–2968

Schindelin J, Arganda-Carreras I, Frise E, Kaynig V, Longair M, Pietzsch T, Preibisch S, Rueden C, Saalfeld S, Schmid B, et al(2012) Fiji: an open-source platform for biological-image analysis. Nat Methods 9: 676–682

Scholthof K-BG, Irigoyen S, Catalan P, Mandadi KK (2018) Brachypodium: A Monocot Grass Model Genus for Plant Biology. Plant Cell 30: 1673–1694

Schopfer P, Oelze-Karow H (1971) [Demonstration of a threshold regulation by phytochrome in the photomodulation of longitudinal growth of the hypocotyl of mustard seedlings (Sinapis alba L.)]. Planta 100: 167–180

Singh AK, Ganapathysubramanian B, Sarkar S, Singh A (2018) Deep Learning for Plant Stress Phenotyping: Trends and Future Perspectives. Trends Plant Sci 23: 883–898

Soga K, Yamazaki C, Kamada M, Tanigawa N, Kasahara H, Yano S, Kojo KH, Kutsuna N, Kato T, Hashimoto T, et al(2018) Modification of growth anisotropy and cortical microtubule dynamics in Arabidopsis hypocotyls grown under microgravity conditions in space. Physiol Plant 162: 135–144

Spalding EP, Miller ND (2013) Image analysis is driving a renaissance in growth measurement. Current Opinion in Plant Biology 16: 100–104

Sudre CH, Li W, Vercauteren T, Ourselin S, Jorge Cardoso M (2017) Generalised Dice Overlap as a Deep Learning Loss Function for Highly Unbalanced Segmentations. Deep Learning in Medical Image Analysis and Multimodal Learning for Clinical Decision Support 240–248

Vandenbussche F, Verbelen J-P, Van Der Straeten D (2005) Of light and length: regulation of hypocotyl growth in Arabidopsis. Bioessays 27: 275–284

van der Walt S, Schönberger JL, Nunez-Iglesias J, Boulogne F, Warner JD, Yager N, Gouillart E, Yu T, scikit-image contributors (2014) scikit-image: image processing in Python. PeerJ 2: e453

Wang L, Uilecan IV, Assadi AH, Kozmik CA, Spalding EP (2009) HYPOTrace: image analysis software for measuring hypocotyl growth and shape demonstrated on Arabidopsis seedlings undergoing photomorphogenesis. Plant Physiol 149: 1632–1637

Wildermann A, Drumm H, Schäfer E, Mohr H (1978a) Control by light of hypocotyl growth in de-etiolated mustard seedlings : I. Phytochrome as the only photoreceptor pigment. Planta 141: 211–216

Wildermann A, Drumm H, Schäfer E, Mohr H (1978b) Control by light of hypocotyl growth in de-etiolated mustard seedlings : II. Sensitivity for newly-formed phytochrome after a light to dark transtition. Planta 141: 217–223

Young JC, Liscum E, Hangarter RP (1992) Spectral-dependence of light-inhibited hypocotyl elongation in photomorphogenic mutants of Arabidopsis: evidence for a UV-A photosensor. Planta 188: 106–114

