## Supplemental Data and Figures for "A deep learning-based approach for high-throughput hypocotyl phenotyping"

**Author affiliations:**

### SUPPLEMENTAL DATA:
### Supplemental figures S1-S3
### Supplemental methods 1 and 2

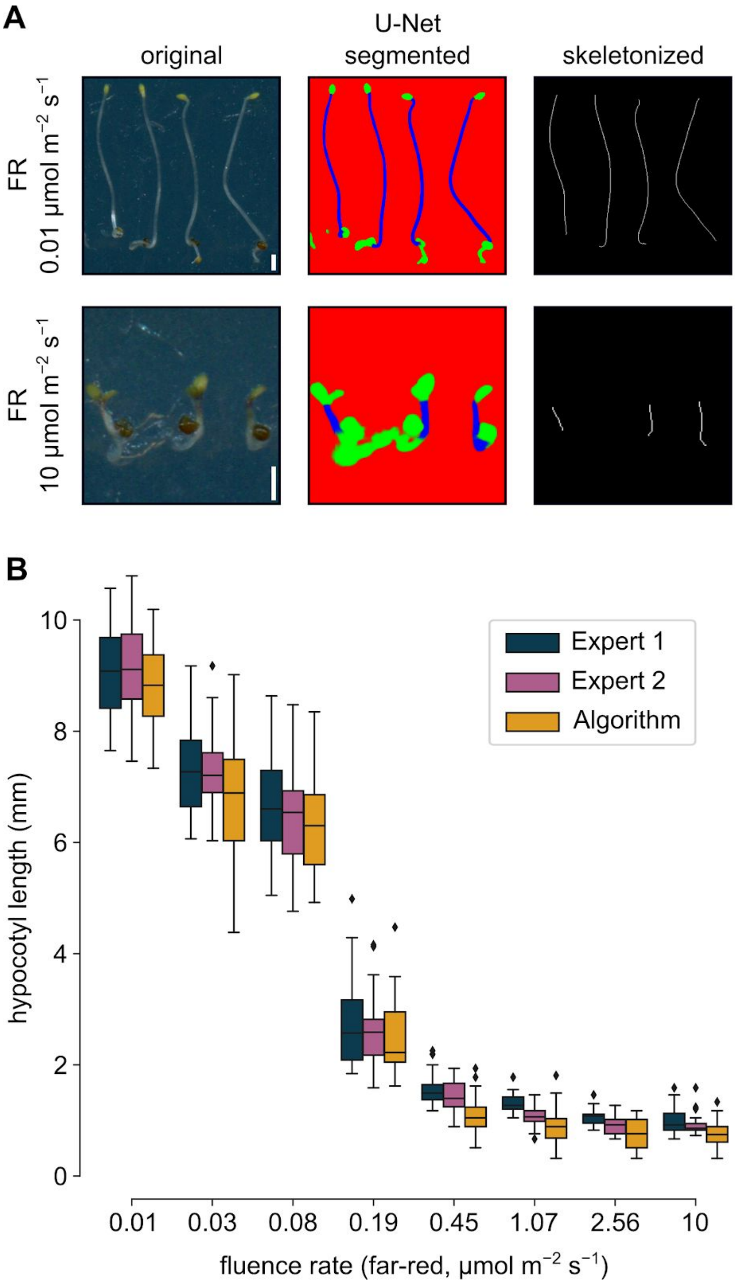


**Supplemental Figure S1. Hypocotyl measurements of Arabidopsis seedlings grown under far-red illumination.**

**(a)** Arabidopsis seedlings were grown on wet filter papers in far-red light for 4 days, placed on agar plate and scanned. A close-up image shows a few seedlings grown under high or low fluences of far-red light, the U-Net segmented and skeletonized image generated from the original by our algorithm. Scale bars represent 1 mm.

**(b)** Fluence rate graph presenting the hypocotyl length values determined by the algorithm and two human experimenters. This box-and-whisker diagram shows the distribution of seedling lengths, where boxes depict the quartiles, whiskers extend to show the rest of the distribution, whereas black diamonds represent outliers. Sample number at every data point is n=30.

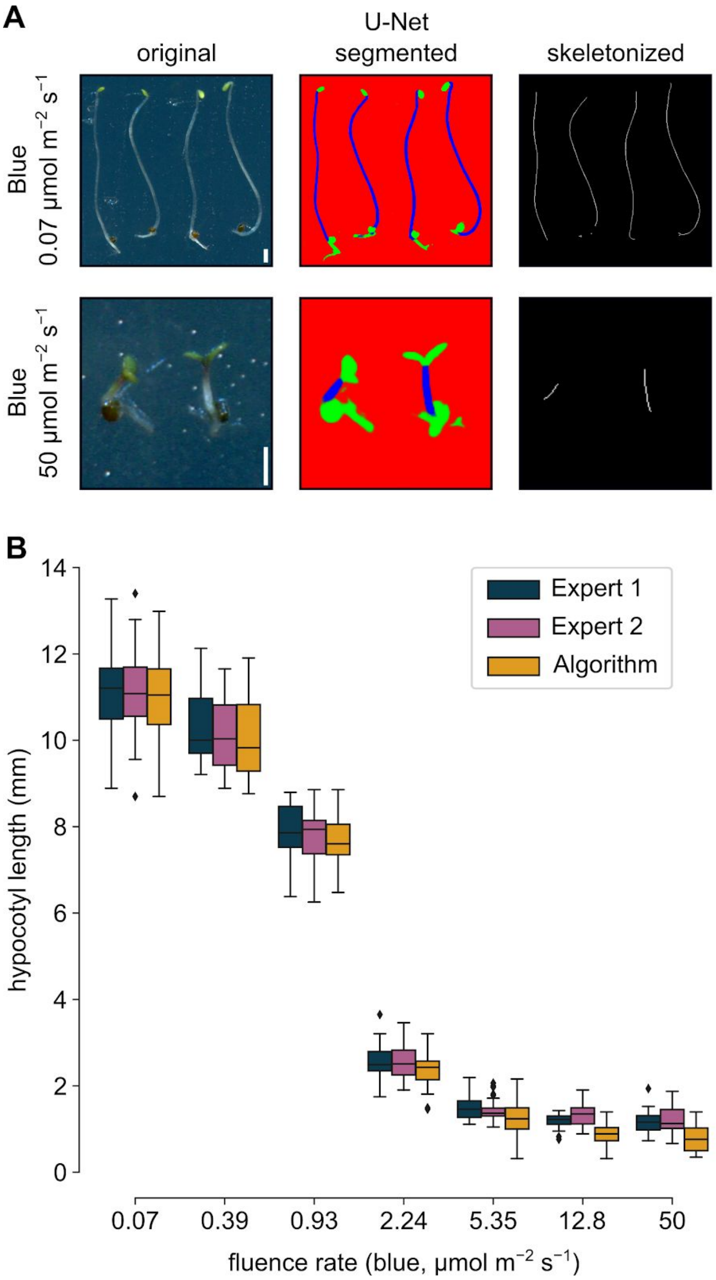


**Supplemental Figure S2. Hypocotyl measurements of Arabidopsis seedlings grown under blue illumination.**

**(a)** Arabidopsis seedlings were grown on wet filter papers in blue light for 4 days, placed on agar plate and scanned. A close-up image shows a few seedlings grown under high or low fluences of blue light, the U-Net segmented and skeletonized image generated from the original by our algorithm. Scale bars represent 1 mm.

**(b)** Fluence rate graph presenting the hypocotyl length values determined by the algorithm and two human experimenters. This box-and-whisker diagram shows the distribution of seedling lengths, where boxes depict the quartiles, whiskers extend to show the rest of the distribution, whereas black diamonds represent outliers. Sample number at every data point is n=30.

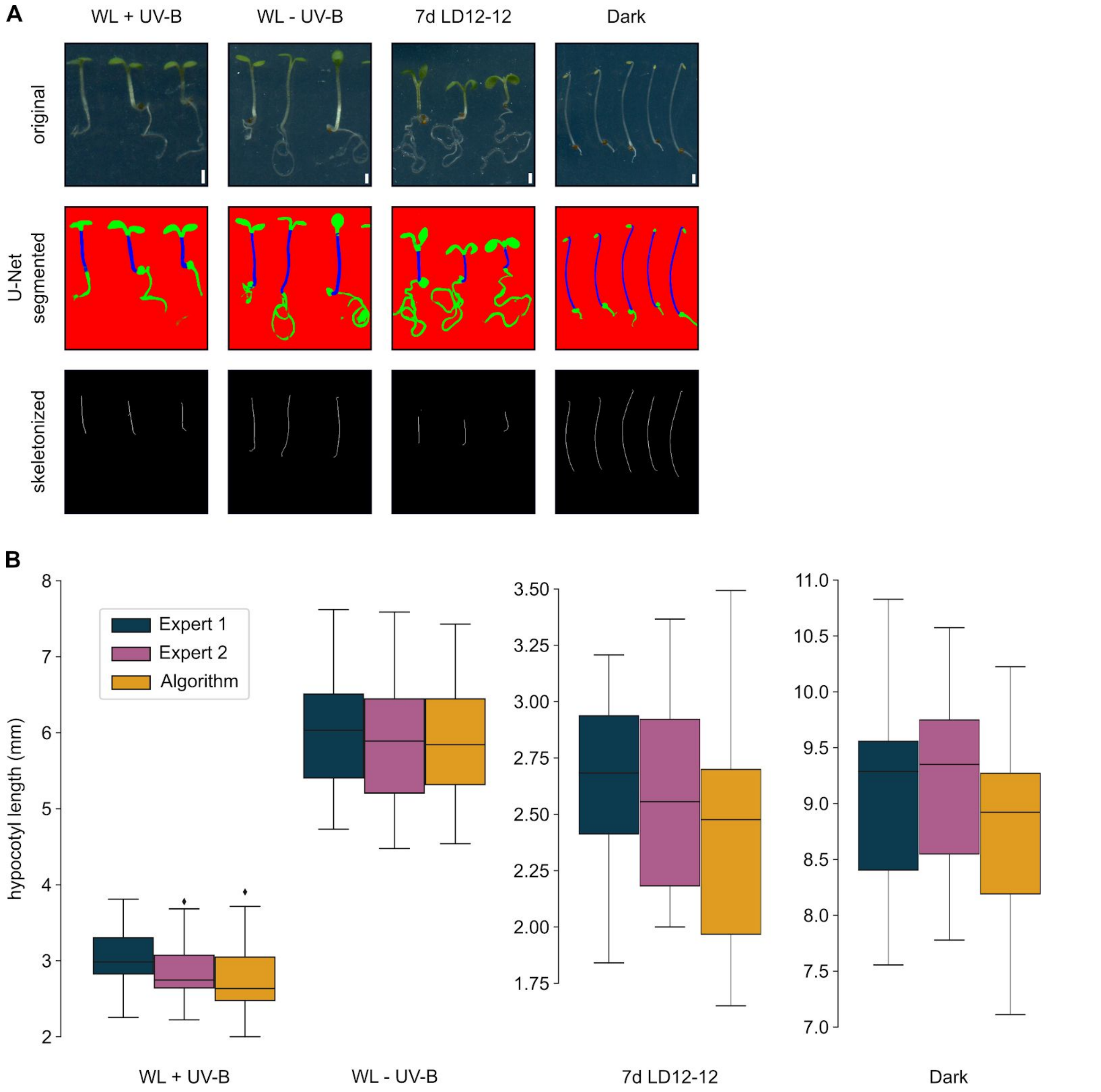


**Supplemental Figure S3. Hypocotyl measurements of Arabidopsis seedlings grown in the dark or under different white light illumination protocols.**

**(a)** Arabidopsis seedlings were grown on wet filter papers in darkness for 4 days (dark), or alternatively on ½ MS plates supplied with 1 % sucrose for 7 days. Seedlings grew for 7 days under 12 h light / 12 h dark photocycles (7d LD 12-12) or were replaced after germination under constant white light supplemented with (WL + UV-B) or without (WL - UV-B) photomorphogenic UV-B irradiation. At the end of the growth phase, seedlings were placed on agar plates and scanned. Close-up images show a few seedlings grown under different conditions detailed above. The U-Net segmented and skeletonized images were generated by our algorithm. Scale bars represent 1 mm.

**(b)** Hypocotyl length values are presented here, which were determined by the algorithm and two human experimenters. This box-and-whisker diagram shows the distribution of seedling lengths, where boxes depict the quartiles, whiskers extend to show the rest of the distribution, whereas black diamonds represent outliers. Sample number at every data point is n=30.

**Supplemental Method 1. Creating custom training data.**
This detailed procedure shows how the U-Net backbone model can be trained on custom data. This process is called annotation, which can be done easily using ImageJ (https://imagej.nih.gov/ij/). During our workflow, we have used the Ubuntu 18.04 operating system (Canonical Ltd., https://ubuntu.com).

**Step 1.** Organize the images to be annotated such that each image is contained in a separate folder with common root. The folder should be named after the image. For example:

```
images_folder
  |-- image_1
      |-- image_1.png
  |-- image_2
      |-- image_2.png
  |-- ...
```

**Step 2.** Open the image to be annotated in ImageJ.

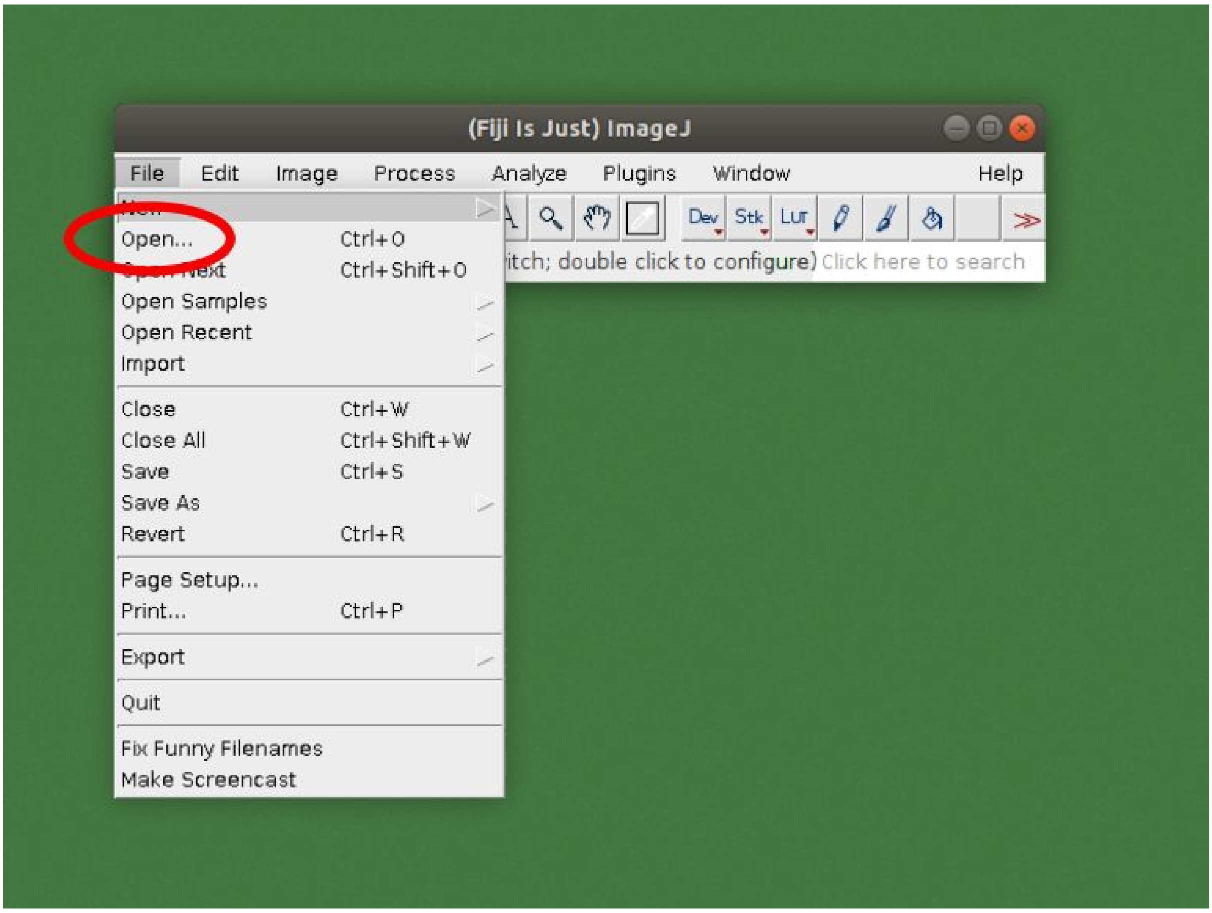

**Step 3.** Open the *ROI manager* tool (ROI - Region Of Interest).

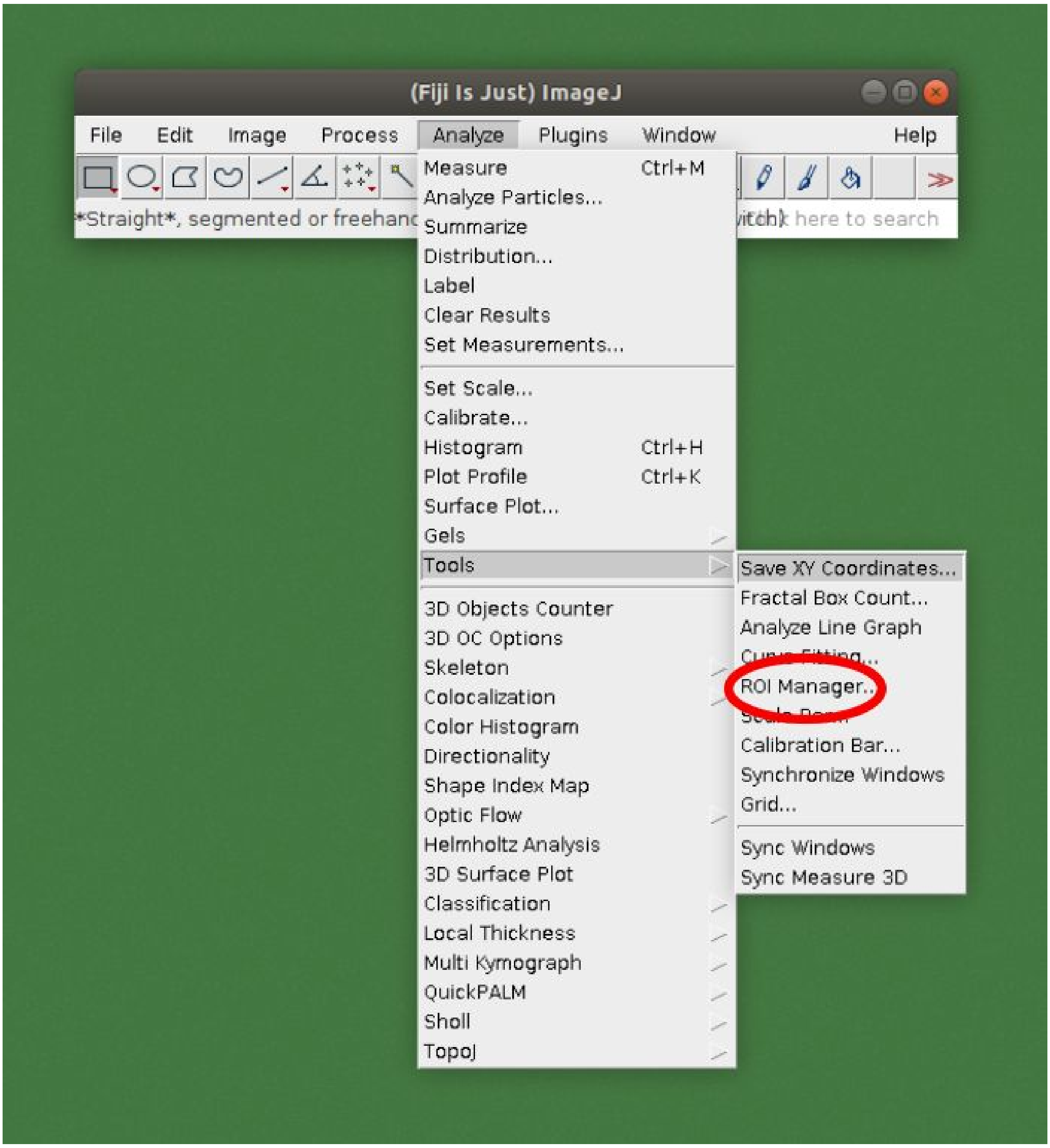

**Step 4**. Select an appropriate selection tool, for example *Freehand selections*.

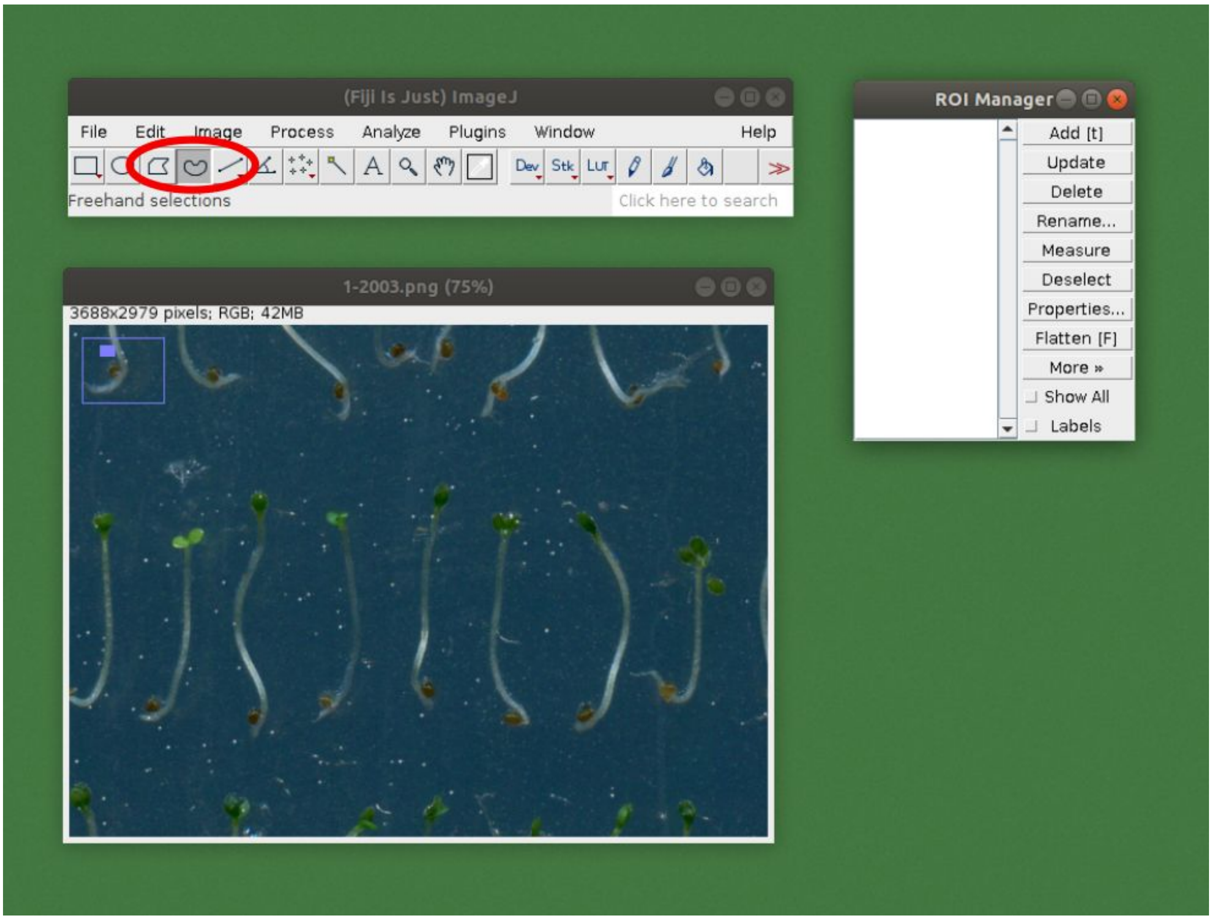

**Step 5.** Outline that part of the plant which should be included in the measurements (hypocotyl in this case). Press *t* or click on the *Add [t]* button to add the selected part to the ROI manager.

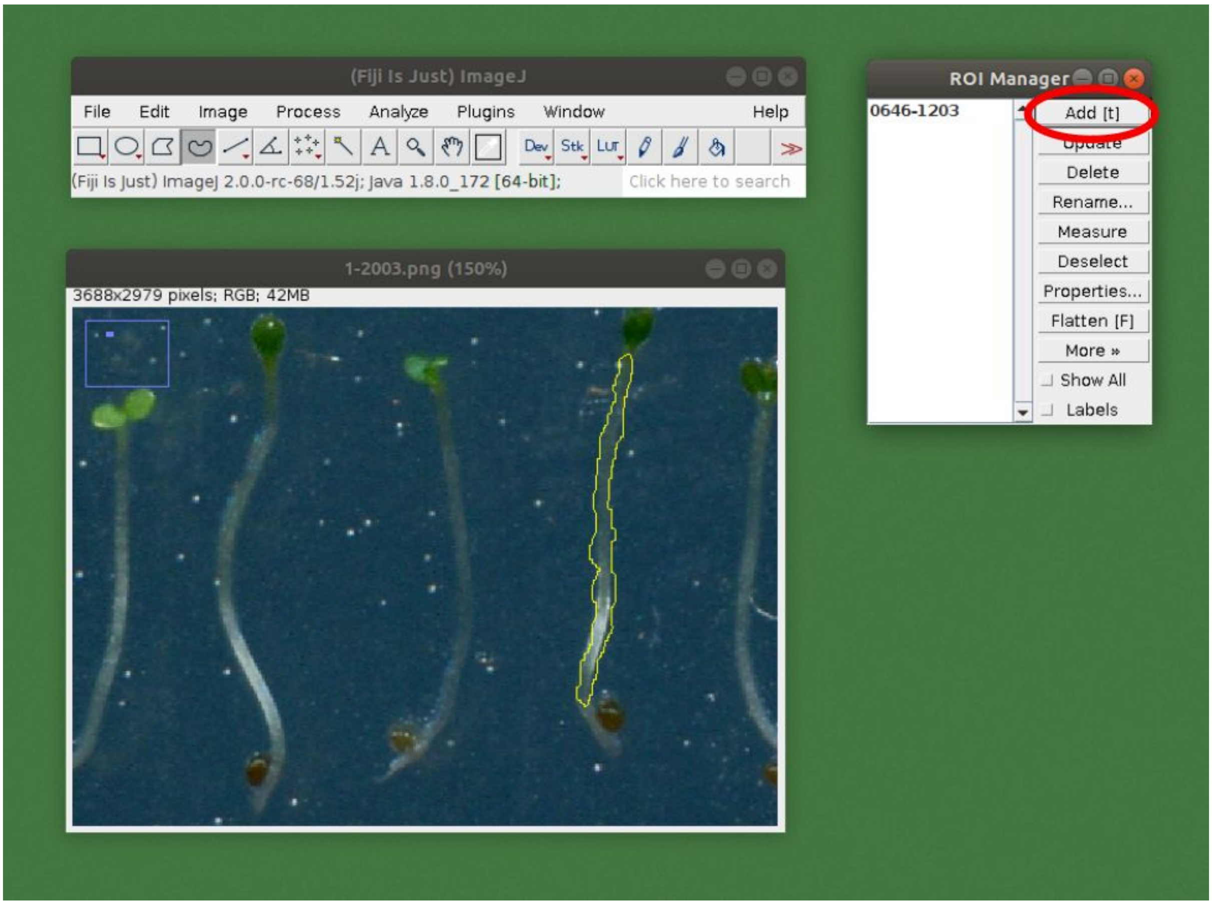

**Step 6.** Repeat the outlining with all seedlings, adding the sections one by one to the ROI manager. WhentThe chosen image is completed, select all ROIs and click *More* > *OR (Combine)*.

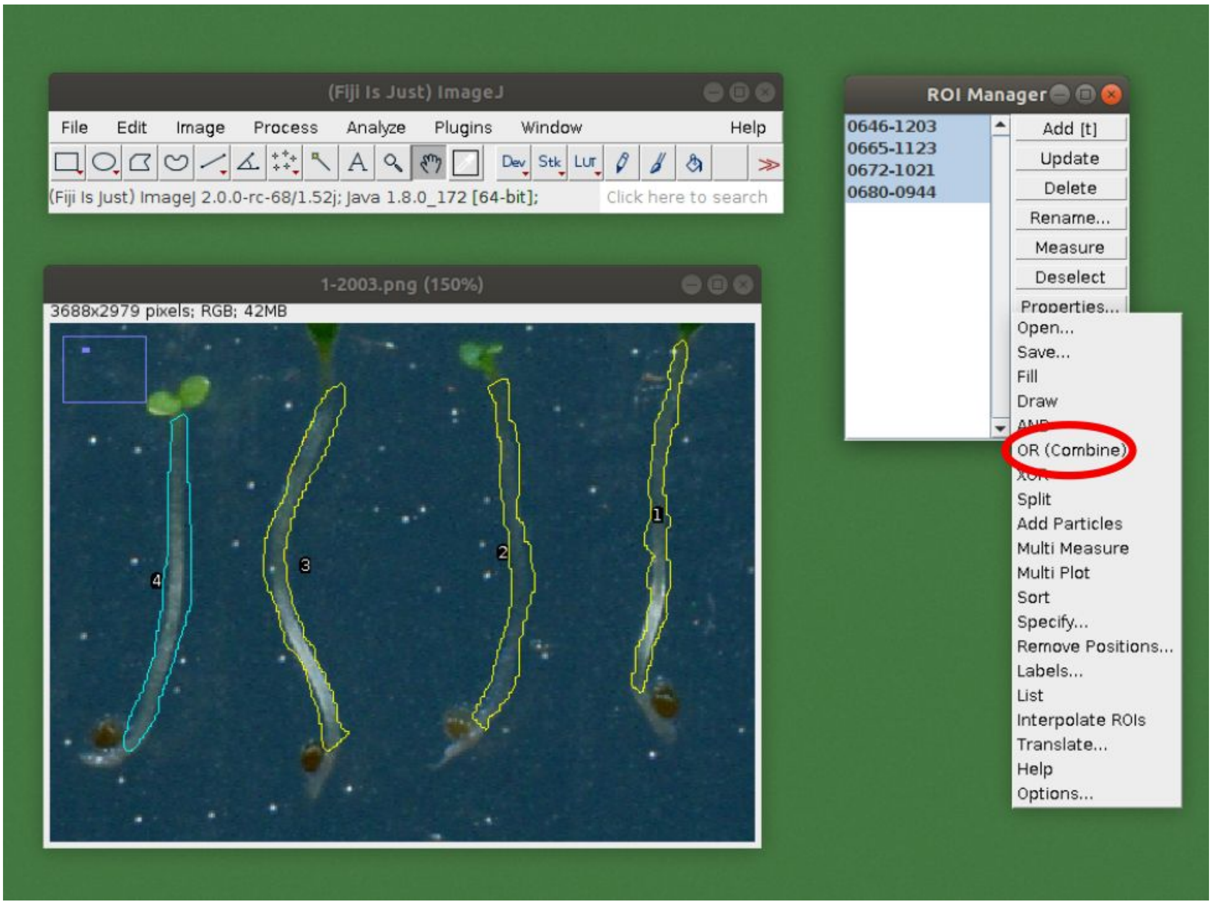

**Step 7.** Press *Edit* > *Selection* > *Create Mask*. This will open up a new image.

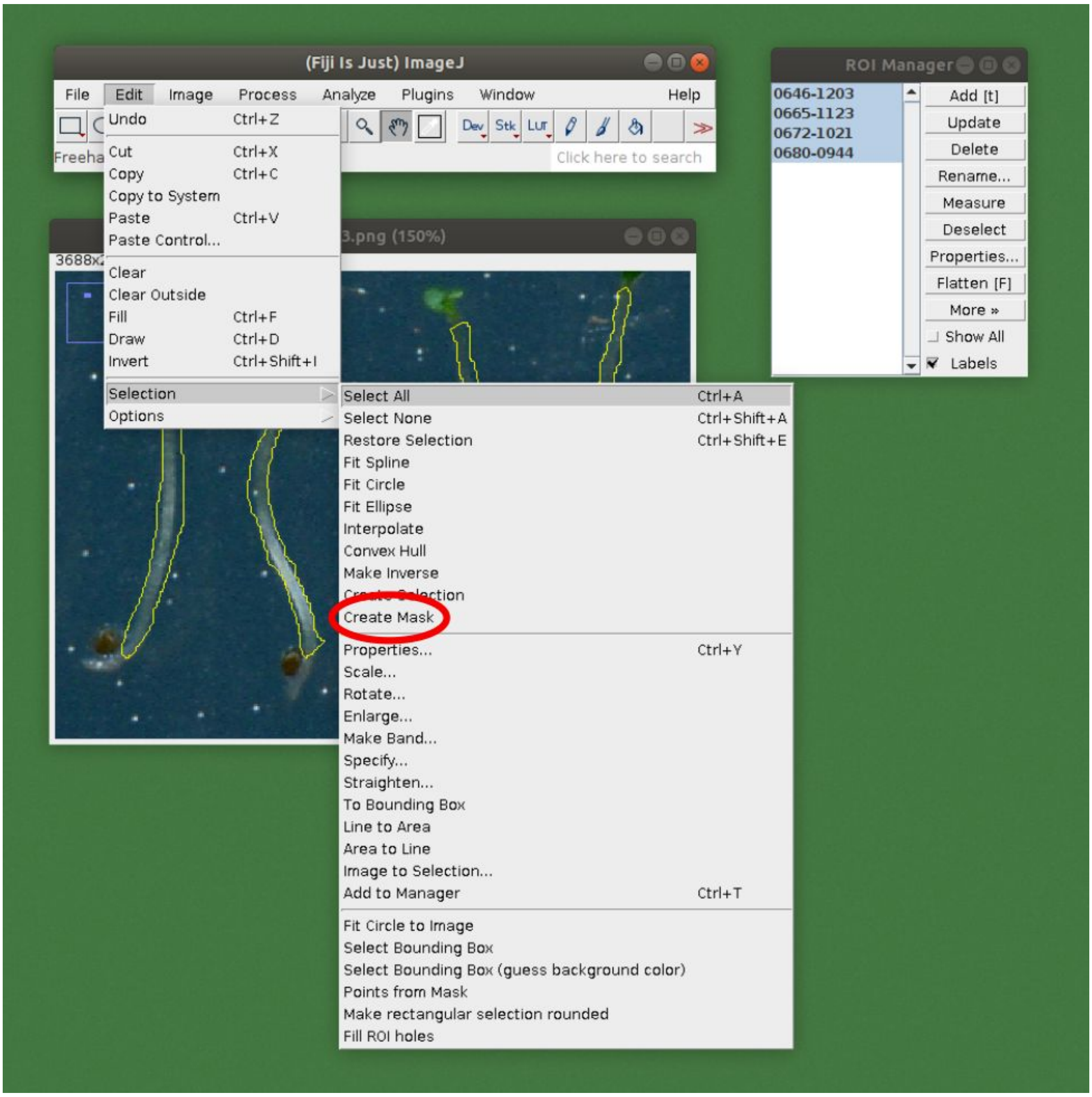

**Step 8.** Invert the mask image by pressing *Edit* > *Invert* or pressing *Ctrl* + *Shift* + *I*.

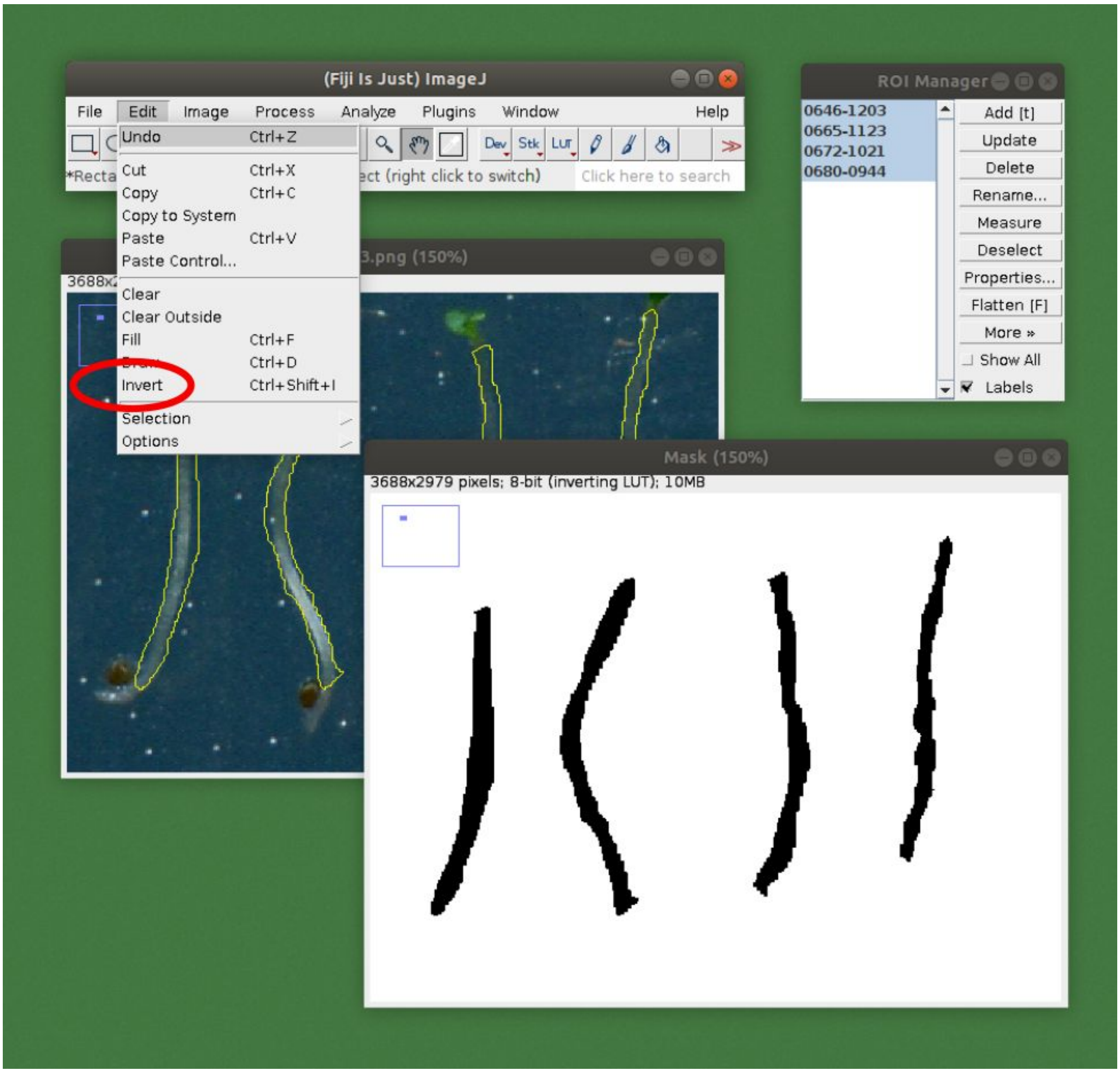

**Step 9.** Save the mask image to the folder along with the original file. Add the suffix -*hypo* to the name. For example, if the original name was *image_1.png*, this image should be named *image_1-hypo.png*.

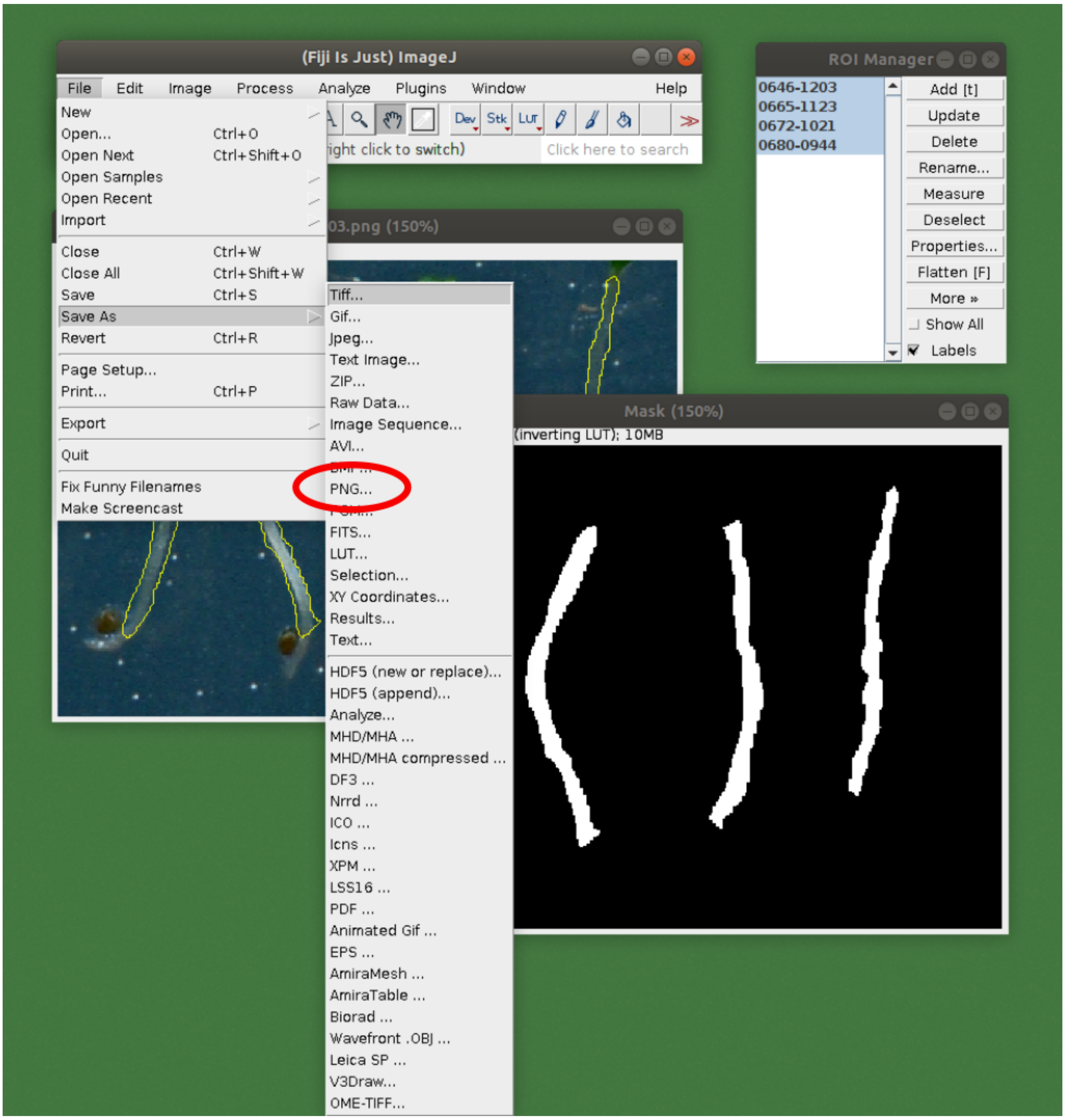

**Step 10.** Repeat the same process to annotate those parts of the plant which should not be included in the measurements. These are the cotyledons and roots (non-hypocotyl parts) in this case. Save the mask again to the same folder. Add the suffix *-nonhypo* to the original file name.

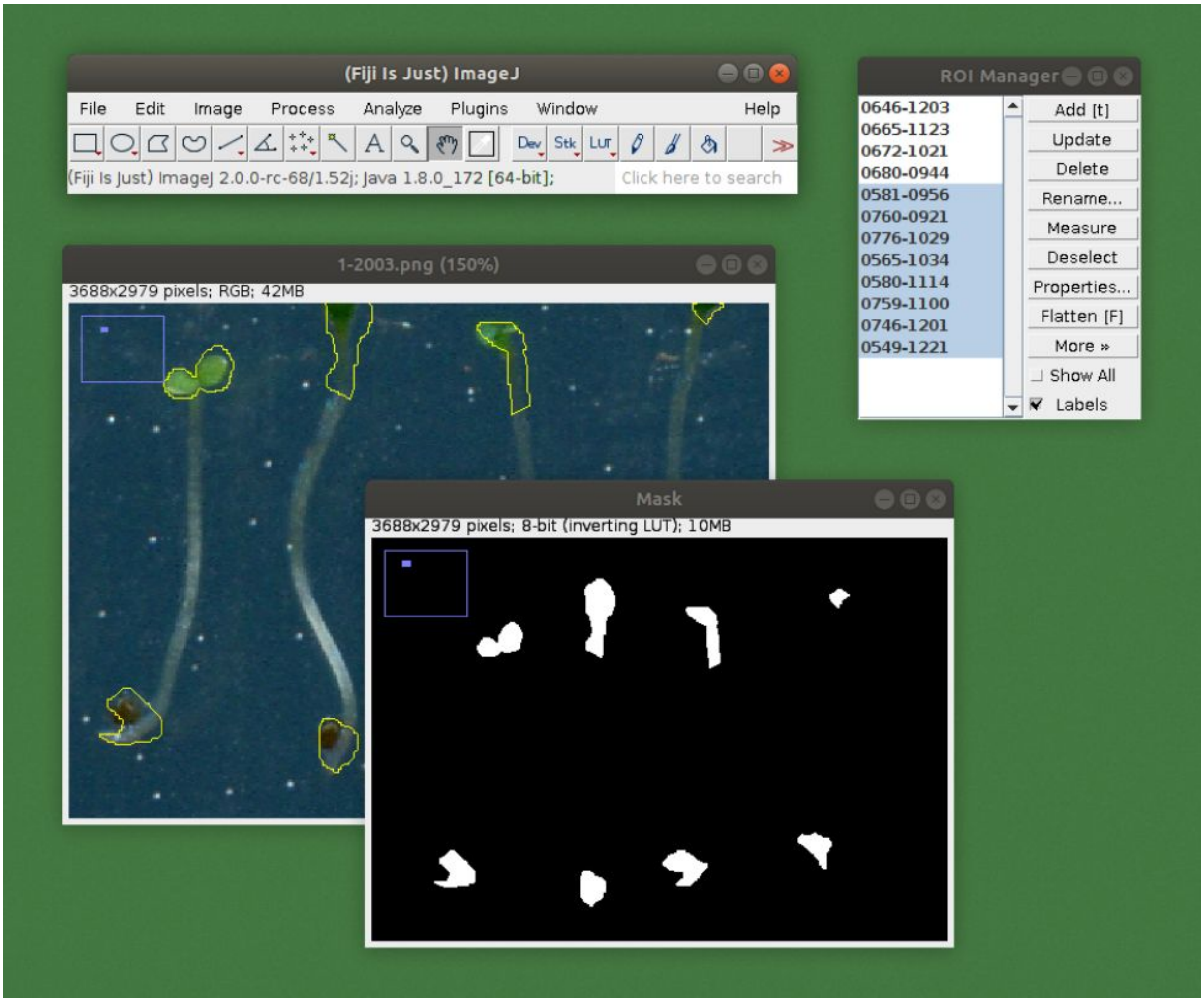


**Step 11.** Create the training data for the algorithm by running the *src/preprocessing/mask.py* in the provided code repository ([https://github.com/biomag-lab/hypocotyl-UNet](https://github.com/biomag-lab/hypocotyl-UNet)). The required arguments are

- `--images_folder`: path to the folder where the folders containing the images and masks located.
- `--export_folder`: path to the folder where the results should be exported.
- `--make_patches`: True if images and masks are to be patched up to smaller pieces. True is recommended for training.

**Supplemental Method 2. Training and using the algorithm.**

The code implementing the method can be found at https://github.com/biomag-lab/hypocotyl-UNet. The following instructions are provided for Linux-based operating systems (we used Ubuntu 18.04).

To use the hypocotyl segmentation tool, clone the repository to the local machine:

```
git clone https://github.com/biomag-lab/hypocotyl-UNet
```

To run the scripts, it is required to have

- Python >= 3.5
- PyTorch >= 0.4
- NumPy >= 1.13
- Pandas >= 0.23
- scikit-image >= 0.14
- matplotlib >= 3.0

The script *src/measure.py* can be used for applying the measuring algorithm on custom images, while *src/train.py* are for training the U-Net model on custom annotated data.

**Step 1.** Training the algorithm. If custom annotated data is available, the containing folder should be organized into the following directory structure:

```
images_folder
  |-- images
      |-- img001.png
      |-- img002.png
      |-- ...
  |-- masks
      |-- img001.png
      |-- img002.png
      |-- ...
```

The mask images should have identical name to their corresponding image. After the training data is organized, the *src/train.py* script can be used to train a custom U-Net model. The required argument is
- `--train_dataset`: path to the folder where the training data is located.
This should match with the `--export_folder` argument given to the *src/preprocessing/mask.py* script during Step 11. of Supplemental Method 1.

The optional arguments are
- `--epochs`: the number of epochs during training. Default is 1000, but this is very dependent on the dataset and data augmentation method used.
- `--batch_size`: the size of the batch during training. Default is 1. If GPU is used, it is recommended to select batch size as large as GPU memory allows.

- `--initial_lr`: initial learning rate. Default is 1e-3, which proved to be the best for the training dataset used. (Different datasets might have more optimal initial learning rates.)
- `--model_name`: name of the model. Default is *UNet-hypocotyl*.
- `--trained_model_path`: to continue training of a previously trained model, its path can be given.
- `--val_dataset`: path to the folder where the validation data is located. During training, validation data is used to catch overfitting and apply early stopping.
- `--model_save_freq`: frequency of model saving. Default is 200, which means that the model is saved after every 200th epoch.
- `--device`: device to be used for the U-Net training. Default is *cpu*, but if a GPU with the CUDA framework installed is available, `cuda:$ID` can be used, where `$ID` is the ID of the GPU. For example, `cuda:0`. (For PyTorch users: this argument is passed directly to the `torch.Tensor.device` object during initialization, which will be used for the rest of the workflow.)

**Step 2.** Using the trained algorithm for prediction. To apply the algorithm on custom images, the folder containing the images should be organized into the following directory structure:

```
images_folder
  |-- images
      |-- img001.png
      |-- img002.png
      |-- ...
```

The *src/measure.py* script can be used to run the algorithm. The required arguments are
- `--images_path`: path to the images folder, which must have the structure outlined above.
- `--model`: path to the U-Net model used in the algorithm. Trained models are available in the *models* folder after training.
- `--result_folder`: path to the folder where results will be exported.

Additionally, you can specify the following:
- `--device`: device to be used for the U-Net training. Default is *cpu*, but if a GPU with the CUDA framework installed is available, `cuda:$ID` can be used, where `$ID` is the ID of the GPU. For example, `cuda:0`. (For PyTorch users: this argument is passed directly to the `torch.Tensor.device` object during initialization, which will be used for the rest of the workflow.)
- `--min_object_size`: the expected minimum object size in pixels. Default is 50. Detected objects below this size will be filtered out.
- `--max_object_size`: the expected maximum object size in pixels. Default is `np.inf`. Detected objects above this size will be filtered out.

For instance, an example is the following:

```
python3 measure.py --images_path path_to_images \
   --model ../models/unet \
   --result_folder path_to_results \
   --device cuda:0
```
